# Altering the redox status of *Chlamydia trachomatis* directly impacts its developmental cycle progression

**DOI:** 10.1101/2024.04.26.591247

**Authors:** Vandana Singh, Scot P. Ouellette

## Abstract

*Chlamydia trachomatis* is an obligate intracellular bacterial pathogen with a unique developmental cycle. It differentiates between two functional and morphological forms: elementary body (EB) and reticulate body (RB). The signals that trigger differentiation from one form to the other are unknown. EBs and RBs have distinctive characteristics that distinguish them, including their size, infectivity, proteome, and transcriptome. Intriguingly, they also differ in their overall redox status as EBs are oxidized and RBs are reduced. We hypothesize that alterations in redox may serve as a trigger for secondary differentiation. To test this, we examined the function of the primary antioxidant enzyme alkyl hydroperoxide reductase subunit C (AhpC), a well-known member of the peroxiredoxins family, in chlamydial growth and development. Based on our hypothesis, we predicted that altering the expression of *ahpC* would modulate chlamydial redox status and trigger earlier or delayed secondary differentiation. To test this, we created *ahpC* overexpression and knockdown strains. During *ahpC* knockdown, ROS levels were elevated, and the bacteria were sensitive to a broad set of peroxide stresses. Interestingly, we observed increased expression of EB-associated genes and concurrent higher production of EBs at an earlier time in the developmental cycle, indicating earlier secondary differentiation occurs under elevated oxidation conditions. In contrast, overexpression of AhpC created a resistant phenotype against oxidizing agents and delayed secondary differentiation. Together, these results indicate that redox potential is a critical factor in developmental cycle progression. For the first time, our study provides a mechanism of chlamydial secondary differentiation dependent on redox status.

## INTRODUCTION

All organisms that are exposed to oxygen are necessarily subjected to oxidative stress. Specifically, the process of metabolizing substrates in the presence of oxygen can generate reactive oxygen species (ROS), which are toxic at high enough concentrations. Thus, from bacteria to humans, systems have evolved to mitigate the accumulation of ROS. At the same time, host defense mechanisms have evolved to leverage ROS production as a means of limiting pathogen growth and survival. Not surprisingly, pathogens have co-evolved to resist these defense mechanisms. For example, many pathogens possess a variety of antioxidant enzymes, such as catalases, glutathione peroxidases, and peroxiredoxins, that help them both subvert ROS-mediated immune system assaults and mitigate metabolic ROS byproducts (Staerck et al., 2017; Wan et al., 2021).

*Chlamydia* is an obligate intracellular bacterium that has significantly reduced its genome size and content in adapting to obligate host dependence. *Chlamydia trachomatis*, the leading cause of bacterial sexually transmitted diseases and preventable infectious blindness, lacks homologs to catalases or glutathione peroxidases but does possess a homolog of Alkyl hydroperoxide reductase subunit C (AhpC). AhpC is a 2-cys peroxiredoxin and is widely conserved in prokaryotes (de Oliveira et al., 2021). Peroxiredoxins can scavenge hydrogen peroxide, peroxynitrite, and organic hydroperoxides (Parsonage et al., 2008; Poole & Ellis, 1996; Seaver & Imlay, 2001) and act as the primary scavenger in pathogens that lack both catalase and glutathione peroxidases (Mastronicola et al., 2014; Richard et al., 2011). Several studies from other bacterial systems have shown that AhpC has a significant role in ROS and RNI scavenging, virulence, and persistence (Cosgrove et al., 2007; Kimura et al., 2012; Oh & Jeon, 2014), and deletion of AhpC results in elevated level of ROS within the bacterium (Zhang et al., 2019). Though *Chlamydia* is dependent on its host for most of its energy requirements, it has some metabolic activities, such as a partial TCA cycle and oxidative phosphorylation (Gerard et al., 2002; Iliffe-Lee & McClarty, 1999), which can be a possible source of intracellular ROS, to which the bacteria must adapt. There is a paucity of knowledge on how *C. trachomatis* modulates oxidative stress. However, some studies explored the effects of redox changes on chlamydial growth. Boncompain et al. reported that infection of *C. trachomatis* induced the transient production of ROS by the host cell at a moderate level for the initial few hours of infection only (Boncompain et al., 2010). Another study also found a similar observation about ROS during infection, reporting that *Chlamydia* requires host-derived ROS for its growth (Abdul-Sater et al., 2010). However, the key mechanisms *Chlamydia* employs to manage ROS-mediated stress have not been characterized.

*Chlamydia* undergoes a complex developmental cycle that comprises two distinct morphological forms, the elementary body (EB) and reticulate body (RB) (Abdelrahman & Belland, 2005). The EB is the smaller (∼0.3 µm), infectious, and nondividing form capable of infecting susceptible host cells. Once internalized into a host-derived vacuole termed an inclusion, the EB differentiates into the non-infectious, larger (∼1 µm), and replicating RB - this process is known as primary differentiation and represents the early phase of the developmental cycle (Clifton et al., 2005). RBs replicate within the inclusion in the midcycle phase using an asymmetric, MreB-dependent polarized division process (Abdelrahman et al., 2016; Lee et al., 2020; Ouellette et al., 2022; Ouellette et al., 2020). In the late phase of the developmental cycle, RBs asynchronously condense into EBs, and this process is termed secondary differentiation. Despite these well-defined differences between chlamydial morphological forms, how *Chlamydia* mechanistically differentiates between functional forms remains unclear.

Intriguingly, a recent study evaluated the redox potential of *Chlamydia* and demonstrated that RBs are reduced, whereas EBs are oxidized (Wang et al., 2014) (Fig. 1A). This is consistent with earlier studies that revealed differences in the crosslinking of outer membrane proteins and type III secretion-related proteins in chlamydial developmental forms (Betts-Hampikian & Fields, 2011; Caldwell et al., 1981; Everett & Hatch, 1995; Wang et al., 2014). Taken together, these observations indicate that, during the developmental cycle, the redox potential of *Chlamydia* is changing. However, whether redox changes in *Chlamydia* directly affect developmental cycle progression is not characterized.

**Fig. 1.**
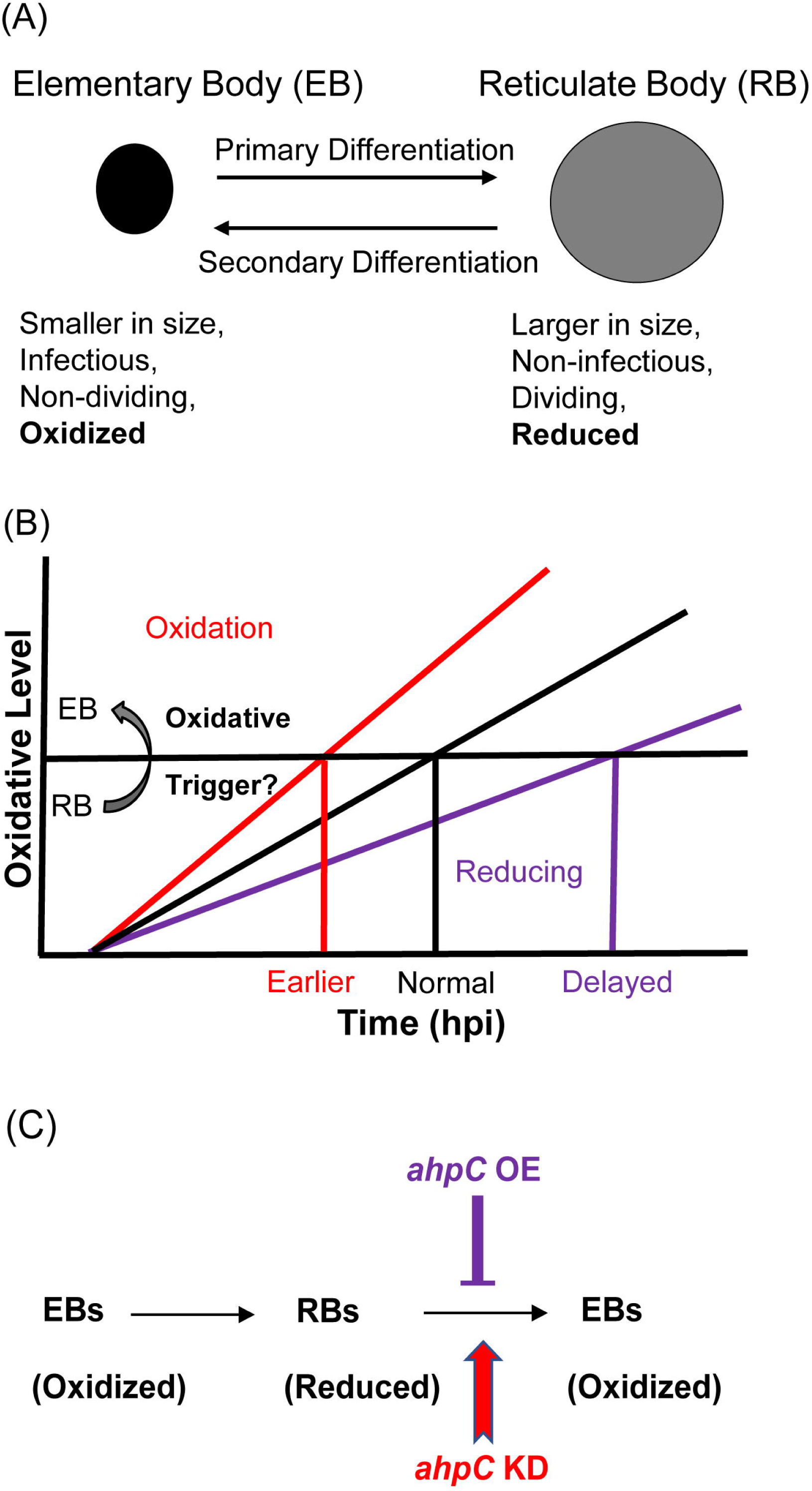
Alterations in redox status of key proteins regulate and drive chlamydial differentiation. (A) Key characteristics of chlamydial developmental forms. (B) Hypothetical model for triggering secondary differentiation through oxidative stress (black angled line). Increasing oxidation of critical protein(s) may lead to earlier differentiation whereas maintaining a reducing environment may delay differentiation. (C) Schematic representation of experimental model for triggering secondary differentiation through altered activity of AhpC. *ahpC* knockdown may lead to earlier differentiation while overexpression of *ahpC* may delay differentiation.

Based on the difference between redox status of EBs and RBs, we hypothesized that changing redox conditions is a critical factor in the process of differentiation from one form to the other. We named this the “redox threshold hypothesis”. In this scenario, as soon as a given RB has crossed an oxidative threshold, the activity of critical proteins is modified to trigger differentiation to the EB (Fig. 1B). We used chlamydial transformants designed to overexpress or reduce AhpC levels to explore the effects of altered redox potential in chlamydial growth and development to test this hypothesis (Fig. 1C). This study establishes the role of AhpC as an antioxidant in *Chlamydia,* as demonstrated by its ability to counteract different peroxide stresses when overexpressed. Overexpression of AhpC had no negative effect on bacterial replication but delayed the differentiation of RBs to EBs. In contrast, under conditions of *ahpC* knockdown, the organism was highly sensitive to oxidizing conditions. Interestingly, this change in redox potential caused earlier expression of EB-associated (late) genes and production of EBs, leading to a shift in developmental cycle progression. This earlier activation of gene expression related to secondary differentiation in *ahpC* knockdown was also observed when developmental cycle progression was blocked by penicillin treatment. Taken together these data provide mechanistic insight into chlamydial secondary differentiation and are the first to demonstrate redox-regulated differentiation in *Chlamydia*.

## RESULTS

### Overexpression of *ahpC* has minimal impact on chlamydial growth

To study the effect of AhpC on chlamydial developmental cycle progression, we first generated an *ahpC* overexpression (OE) strain using a plasmid encoding an anhydrotetracycline (aTc) inducible *ahpC* (untagged) and transformed it into a *C. trachomatis* L2 strain lacking its endogenous plasmid (-pL2). The same plasmid vector backbone (i.e., encoding mCherry in place of *ahpC*) was used as an empty vector control (EV). HeLa cells were infected with these transformants, and, at 10 hpi, expression of the construct was induced or not. Lacking an antibody against AhpC, overexpression of *ahpC* was validated by reverse transcription-quantitative PCR (RT-qPCR). Here, an approximate 1-log increase in transcripts of *ahpC* was detected at 14 and 24 hpi in the induced strain in comparison to the uninduced strain (Fig. 2A). Immunofluorescence analysis (IFA) was performed at 14 and 24 hpi to examine the organisms’ morphology and overall inclusion growth. For IFA, individual bacteria were labeled with an anti-major outer membrane protein (MOMP) antibody, and DAPI was used to stain for DNA. IFA imaging revealed that there was no observable impact on the morphology of the EV control strain under the conditions tested. In contrast, we did note that overexpression of *ahpC* increased the overall inclusion area (Fig. 2B and 2C). We next quantified the total number of bacteria (i.e., both RBs and EBs) by measuring genomic DNA and observed a significant increase in gDNA levels at 24 hpi in response to increased *ahpC* expression (Fig. 2D). Although inclusions were larger in area and contained more total bacteria as assessed by gDNA levels when overexpressing *ahpC*, recoverable inclusion forming units (IFUs: a measure of infectious EBs) were significantly lower under these conditions at 24 hpi (Fig. 2E). However, by 48 hpi, the difference between the induced and uninduced conditions, although still reduced, was not statistically significant. The average number of IFUs was 5.30×10^7^ and 1.56×10^9^ for the *ahpC* OE strain in the uninduced condition at 24 and 48 hpi, respectively. For the EV strain, the average number of IFUs was 7.07×10^6^ and 1.90×10^8^ in the uninduced condition at 24 and 48 hpi, respectively. The inclusion forming unit analysis only measures viable EBs from a population, suggesting that AhpC overexpression skews the ratio of RBs to EBs. To test this, we calculated the ratio of IFUs to gDNA at 24 hpi, which revealed that the relative number of RBs is higher (and EBs lower) as a result of overexpression of *ahpC* (Fig 2E). Overall, these data support the hypothesis that overexpressing *ahpC* delays production of infectious EBs.

**Fig. 2.**
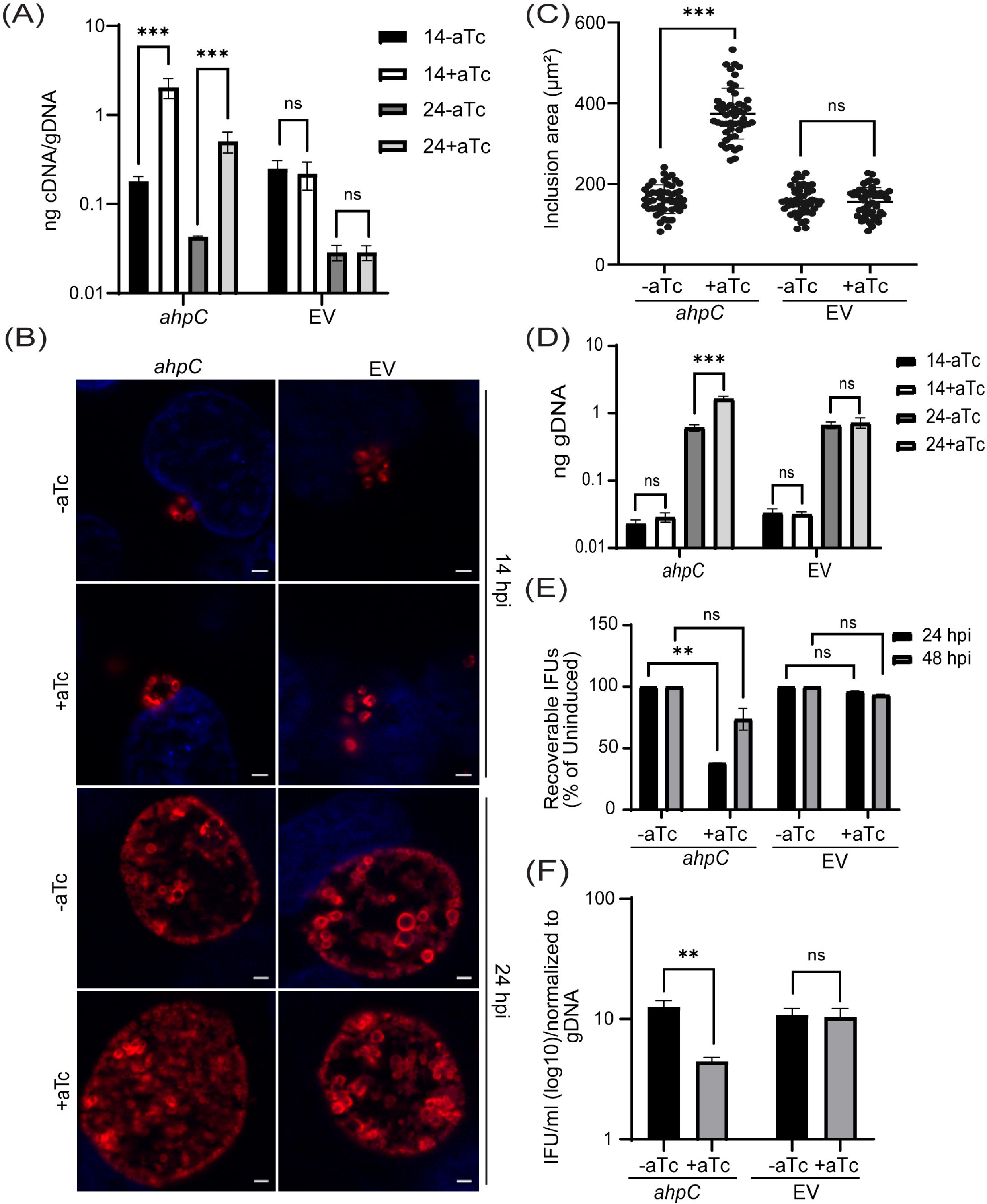
Overexpression of *ahpC* affects chlamydial growth and differentiation. (A) Transcriptional analysis of *ahpC* in *ahpC* overexpression (*ahpC*) and empty vector (EV) control using RT-qPCR following induction at 10 hpi with 1 nM aTc. RNA and genomic DNA (gDNA) were harvested at 14 and 24 hpi and processed as mentioned in materials and methods. Data are presented as a ratio of cDNA to gDNA plotted on a log scale. \*\*\**p*< 0.0001 vs uninduced sample by using two-way ANOVA. Data represent three biological replicates. (B) Immunofluorescence assay (IFA) of *ahpC* and EV at 14 and 24 hpi. Construct expression was induced or not at 10 hpi with 1 nM aTc, and samples were fixed with methanol at 24 hpi then stained for major outer membrane protein (MOMP - red) and DAPI (blue) to label DNA. Scale bars = 2 µm. Images were captured using a Zeiss Axio Imager Z.2 with Apotome2 at 100x magnification. Representative images of three biological replicates are shown. (C) Impact of *ahpC* overexpression on inclusion area. Inclusion area of *ahpC* overexpression and EV strains was measured using ImageJ. Experimental conditions were the same as mentioned in section (B). The area of 50 inclusions was measured per condition for each sample. \*\*\**p*< 0.001 vs uninduced sample by using ordinary one-way ANOVA. Data were collected from three biological replicates. (D) IFU assay of *ahpC* overexpression and empty vector control. Expression of the construct was induced or not at 10 hpi, and samples were harvested at 24 or 48 hpi for reinfection and enumeration. IFUs were calculated as the percentage of uninduced samples. \*\**p*< 0.001 vs uninduced sample by using multiple paired t test. Data represent three biological replicates. (E) Quantification of genomic DNA (gDNA) determined by qPCR in *ahpC* overexpression and empty vector control. Construct expression was induced or not at 10 hpi with 1 nM aTc, gDNA was harvested at 14 and 24 hpi, and ng gDNA were plotted on a log scale. \*\*\**p*< 0.0001 vs uninduced sample by using two-way ANOVA. Data represent three biological replicates. (F) Ratio of log10 IFUs and log10 gDNA. IFU/ml from (E) was normalized with gDNA from (D). \*\**p*< 0.001 vs uninduced sample by using multiple unpaired t test. Data represent three biological replicates.

### Overexpression of *ahpC* confers resistance to peroxides in *Chlamydia*

AhpC is an important peroxiredoxin involved in oxidative damage defense. Before determining whether increased *ahpC* expression impacts chlamydial sensitivity to oxidizing agents, we first sought to determine the response of *C. trachomatis* to inorganic or organic hydroperoxides such as hydrogen peroxide (H_2_O_2_), cumene hydroperoxide (CHP), and tert-butyl hydroperoxide (TBHP) as well as peroxynitrite (PN). To do this, we evaluated the sensitivity of HeLa cells, infected or not with the empty vector (EV) control strain, to different concentrations of oxidizing agents using a viability assay. Both uninfected and EV-infected HeLa cells tolerated up to 1 mM oxidizing agents for 30 min (Fig. S1). Next, different concentrations (lower than 1 mM) of these oxidants were tested on wild-type (WT) *C. trachomatis* L2/434/Bu (*Ctr* L2). HeLa cells were infected with *Ctr* L2 and, at 16 hpi, were exposed to different concentrations of exogenous oxidizing agents for 30 min only, before washing out the oxidizing agents and replacing the media. Samples for IFU and IFA assays were then collected at 24 hpi to assess effects of the oxidizing agents on growth of *Ctr* L2. 62.5 µM concentration of all three inorganic and organic peroxides had no appreciable effect on IFUs, and inclusion size and morphology also remained unaffected at this sublethal concentration. The concentrations of inorganic and organic peroxides that resulted in a decrease in IFUs to ∼50% of the untreated culture were 500 µM (H_2_O_2_) and 250 µM (CHP and TBHP), respectively, with a concurrent decrease in inclusion size. 1 mM H_2_O_2_, CHP, and TBHP caused >90% reduction in IFUs and small inclusions. PN at 1 mM concentration was less effective than other oxidizing agents and showed only a ∼30% decline in IFUs compared to the untreated culture (Fig. S2).

To further examine the antioxidant functions of AhpC, we next exploited our *ahpC* overexpression strain to explore its capacity to protect chlamydiae from oxidizing agents. Both *ahpC* OE and EV strains were used to infect HeLa cells and, at 10 hpi, expression of the constructs was induced or not with 1 nM aTc. At 16 hpi, infected cells were exposed to various concentrations of exogenous oxidizing agents for 30 min. At 24 hpi, IFU and IFA samples were collected to measure chlamydial growth and assess chlamydial morphology, respectively. As shown in Figure 3A, 3B, and S3, the *ahpC* overexpression strain showed increased resistance to all oxidants tested as the mean IFUs and the size of inclusions were greater than that of the uninduced but treated control. There was no change in bacterial growth and morphology in the case of 62.5 µM concentrations of oxidizing agents, which was anticipated since we determined this was a sublethal concentration. The resistant phenotype was more evident at the higher concentrations of H_2_O_2_, CHP, and TBHP. Overexpression of *ahpC* also contributed to higher resistance against PN. Conversely, the empty vector control strain behaved as expected (i.e., like WT) in the presence of oxidizing agents, with similar results observed in the presence or absence of aTc (Fig. 3C, 3D, and S3). Collectively, these data demonstrate that the chlamydial AhpC possesses antioxidant activity, as expected, and that increased *ahpC* expression is protective for *Chlamydia* in the presence of increased oxidative stress.

**Fig. 3.**
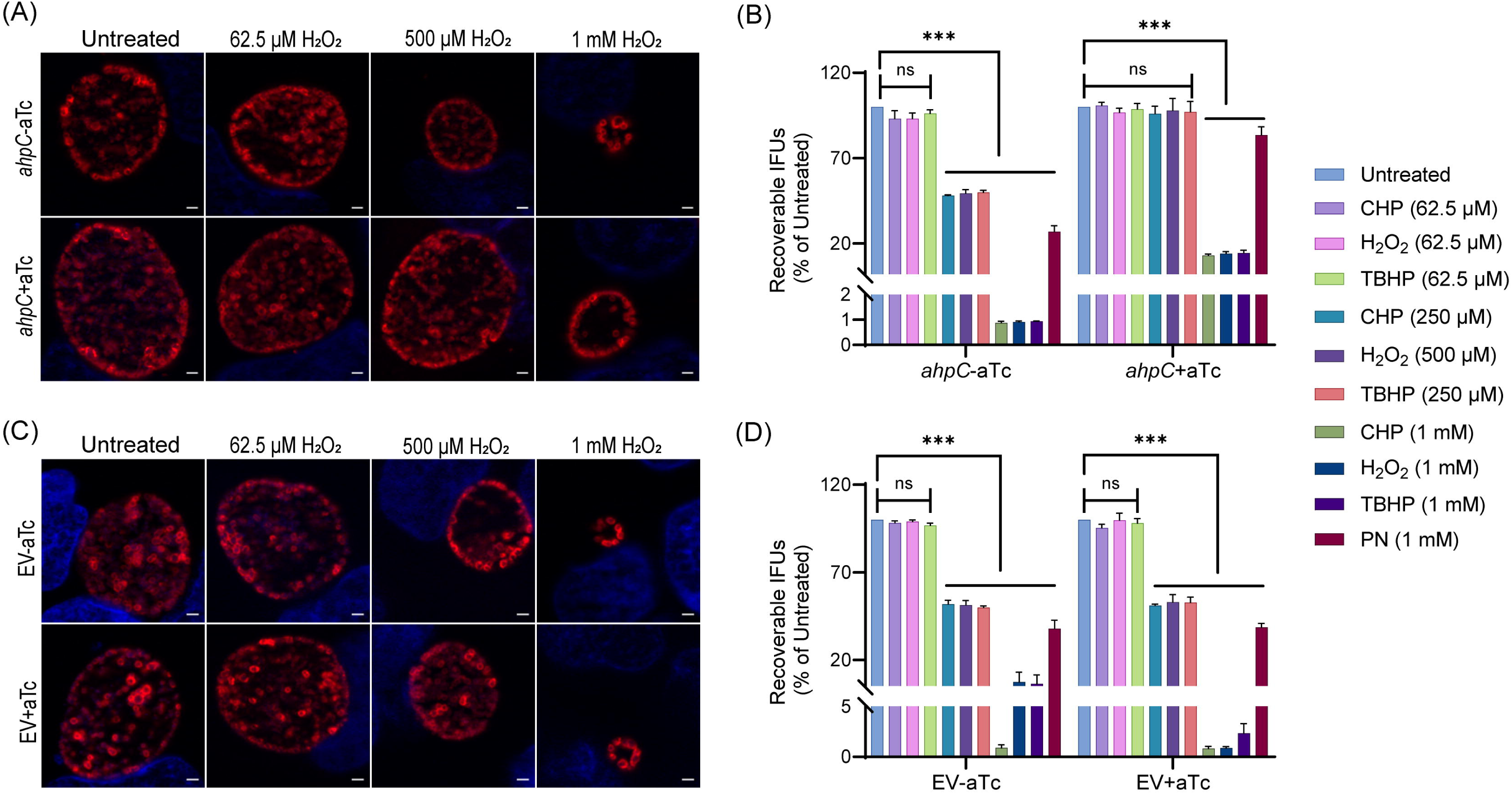
Higher expression of *ahpC* provides resistance to peroxides in *Chlamydia.* IFA of *ahpC* (A) or EV (C) exposed to oxidizing agents. Construct expression was induced or not at 10 hpi with 1 nM aTc, and samples were treated with three different concentrations of hydrogen peroxide (H_2_O_2_) at 16 hpi for 30 min then fixed with methanol at 24 hpi and stained and imaged as described in the legend of Fig. 2B. Representative images from three biological replicates are shown. IFU analysis of *ahpC* (B) or EV (D) following treatment with oxidizing agents, CHP-Cumene hydroperoxide, H_2_O_2_-Hydrogen peroxide, TBHP-Tert-butyl hydroperoxide, and PN-Peroxynitrite. Samples were processed as described for (A) and (C), and IFUs harvested at 24 hpi. IFUs of treated samples were compared with respective untreated controls. \*\*\**p*< 0.0001 vs untreated sample by using two-way ANOVA. Data represent three biological replicates.

### Knockdown of *ahpC* negatively impacts chlamydial growth

To further define the function(s) of AhpC in the chlamydial developmental cycle, we used a novel dCas12-based CRISPR interference (CRISPRi) strategy adapted for *Chlamydia* by our lab (Ouellette et al., 2021). We generated an *ahpC* knockdown strain harboring the pBOMBL12CRia with a crRNA targeting the *ahpC* 5’ intergenic region and overlapping the ATG start site (plasmid designated as pL12CRia(*ahpC*)). We used a strain carrying a pL12CRia plasmid with a crRNA with no homology to any chlamydial sequence (i.e., non-targeting [NT]) to serve as a negative control. To confirm the knockdown of *ahpC*, RT-qPCR was employed. Here, we infected HeLa cells with the *ahpC* knockdown (*ahpC* KD) or NT strains and induced dCas12 expression or not at 10 hpi using 1 nM aTc. Nucleic acid samples were harvested at 14 hpi and 24 hpi. RT-qPCR analysis of the *ahpC* KD revealed approximately 90% reduction of *ahpC* transcripts compared to the uninduced control, thus confirming the knockdown of *ahpC* (Fig. 4A). In the case of the NT control, there was no effect on *ahpC* transcripts. IFA of these strains revealed noticeably smaller inclusions after blocking expression of *ahpC* as compared to the uninduced sample or the NT conditions (Fig. 4B). We next measured total bacterial counts using genomic DNA. Surprisingly, even though inclusions were smaller during *ahpC* knockdown, we observed higher gDNA levels at 14 hpi compared to the uninduced control, followed by only a small increase from 14 to 24 hpi (Fig. 4C). In contrast, the uninduced strain showed a logarithmic increase in gDNA levels during this timeframe, similar to the NT strain (Fig. 4C). To further explore the effect of reduced *ahpC* transcripts in *Chlamydia*, IFU assays were performed. IFU analysis exhibited severely reduced progeny (>90%) in *ahpC* knockdown compared to its respective uninduced control at both time points assessed (24 and 48 hpi) (Fig. 4D). In contrast, the NT strain showed less than a 50% reduction after inducing dCas12 expression at these timepoints consistent with prior observations that dCas12 expression slightly delays developmental cycle progression (Hatch & Ouellette, 2023; Reuter et al., 2023). The ratio of IFUs to gDNA at 24 hpi is significantly higher for the *ahpC* KD, suggesting a lower number of RBs in proportion to EBs (Fig 4E). Taken together, these analyses indicate that reducing *ahpC* levels and/or activity severely reduced chlamydial growth at 24 and 48 hpi.

**Fig. 4.**
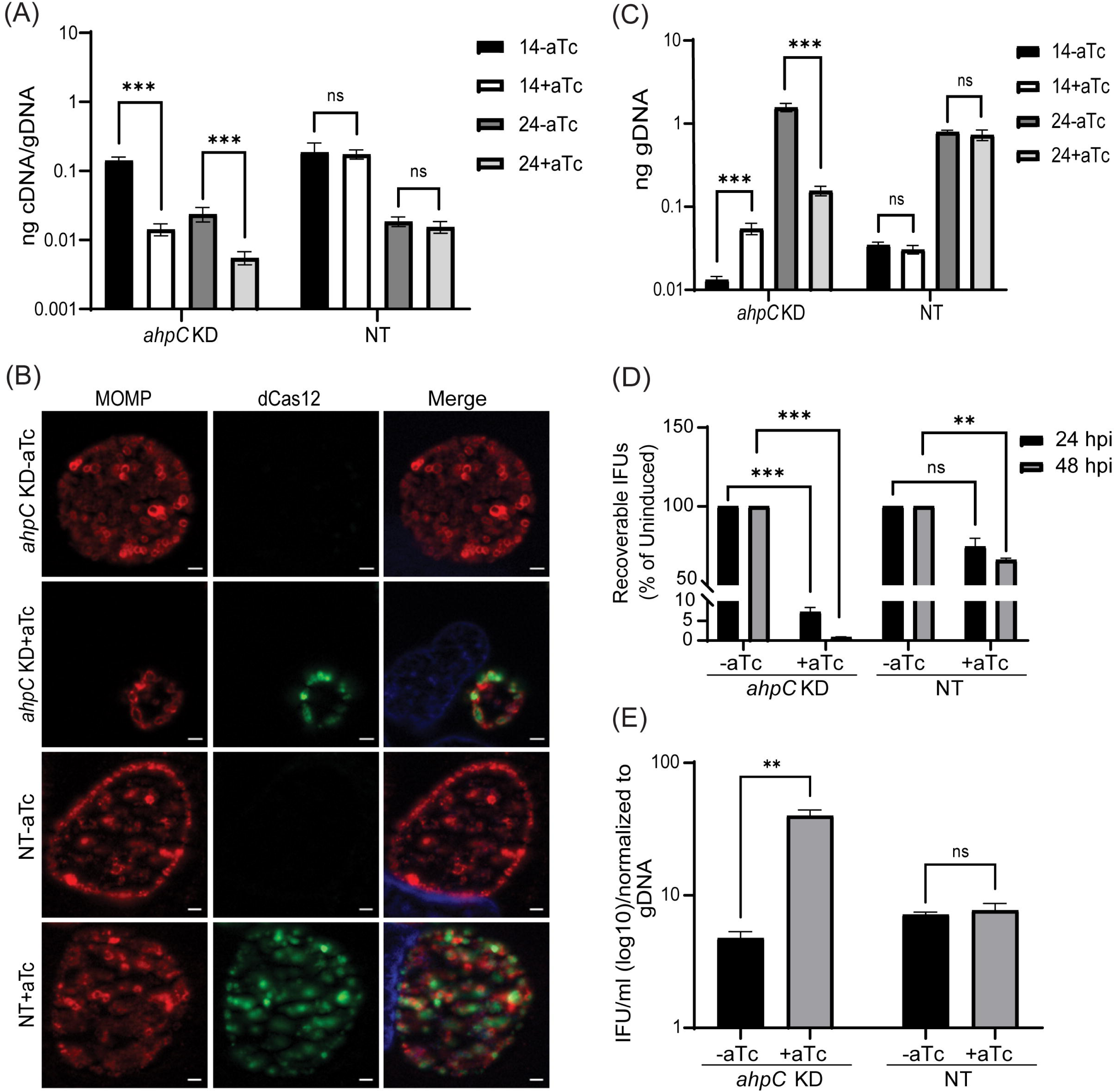
Reduced levels of AhpC negatively impact chlamydial growth. (A) Transcriptional analysis of *ahpC* in knockdown (*ahpC* KD) and non-target (NT) control using RT-qPCR following induction at 10 hpi with 1 nM aTc. RNA and gDNA were harvested at 14 and 24 hpi. Quantified cDNA was normalized to gDNA, and values were plotted on a log scale. \*\*\**p*< 0.0001 vs uninduced sample by using two-way ANOVA. Data represent three biological replicates. (B) IFA was performed to assess inclusion size and morphology using the same induction conditions as in section (A). At 24 hpi, cells were fixed with methanol and stained using primary antibodies to major outer membrane protein (MOMP), Cpf1 (dCas12), and DAPI. All images were acquired on Zeiss Axio Imager Z.2 with Apotome2 at 100x magnification. Bars, 2 µm. Representative images of three biological replicates are shown. (C) Quantification of genomic DNA (gDNA) determined by qPCR in *ahpC* KD and NT strains. dCas12 expression was induced or not at 10 hpi, and gDNA was harvested at 14 and 24 hpi and plotted on a log scale. \*\*\**p*< 0.0001 vs uninduced sample by using two-way ANOVA. Data represent three biological replicates. (D) IFU titers following induction at 10 hpi with 1 nM aTc. IFUs were counted from 24 and 48 hpi samples and calculated as percentage of uninduced samples. \*\*\**p*< 0.0001, \*\**p*< 0.001 vs uninduced sample by using multiple paired t test. Data represent three biological replicates. (E) Ratio of log10 IFUs by log10 gDNA. IFU/ml from (D) was normalized with gDNA from (C). \*\**p*< 0.001 vs uninduced sample by using multiple unpaired t test. Data represent three biological replicates.

To investigate whether *ahpC* knockdown resulted in increased ROS levels in the bacteria (Zhang et al., 2019), the intracellular ROS levels were measured in infected cells in *ahpC* knockdown conditions and compared to the uninduced control condition and uninfected cells. We used the cell-permeable, fluorogenic dye CellROX Deep Red to measure ROS levels. This dye remains non-fluorescent in a reduced state and exhibits bright fluorescence once oxidized by ROS. ROS generation was measured in uninfected and *ahpC* KD-infected HeLa cells at 24 and 48 hpi. As shown in Figure 5A, *ahpC* knockdown resulted in significantly higher ROS than the uninfected or infected but uninduced samples at both 24 hpi and 48 hpi time points. We also performed live-cell microscopy for visualization of ROS generation in uninfected and *ahpC* KD-infected HeLa cells at 24 and 40 hpi. These microscopy images revealed that *ahpC* KD resulted in higher ROS in comparison to the uninduced and uninfected cells (Fig. 5B and S4). Importantly, the signal was associated with chlamydial inclusions and not the host cell, thus indicating that the measurements from Figure 5A were from the bacteria. These results together indicate that AhpC knockdown results in increased ROS levels in *C. trachomatis*.

**Fig. 5.**
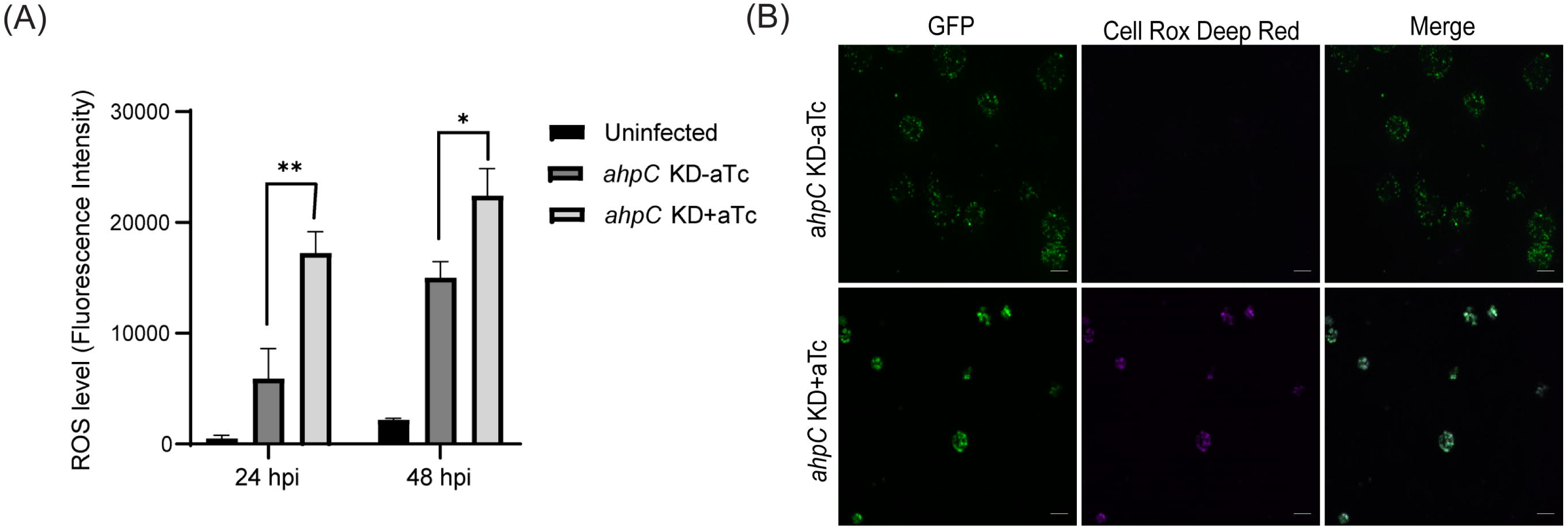
Intracellular ROS levels were measured to investigate the function of AhpC in reducing ROS. (A) HeLa cells were infected or not with *ahpC* knockdown, and construct was induced or not at 10 hpi with 1 nM aTc. At 24 or 48 hpi, samples were washed with DPBS and incubated with CellROX Deep red dye for 30 min in dark. ROS levels were measured at wavelengths of 640 nm (excitation) and 665 nm (emission). \*\**p*< 0.001, \**p*< 0.01 vs uninduced sample by using two-way ANOVA. Data represent three biological replicates. (B) Microscopy images were acquired using live cells on Zeiss Axio Imager Z.2 with Apotome2 at 100x magnification. Scale Bars= 10 µm. Representative images of three biological replicates are shown.

### Knockdown of *ahpC* sensitizes *Chlamydia* to oxidizing agents

As overexpression of *ahpC* resulted in increased resistance to oxidizing agents, we next explored whether knockdown of *ahpC* resulted in increased sensitivity to such agents. As described above, *ahpC* KD and NT strains were used to infect HeLa cells and, at 10 hpi, expression of dCas12 was induced or not with 1 nM aTc. Considering that the *ahpC* KD already demonstrated reduced IFUs and inclusion sizes at 24 hpi, only sublethal concentrations (62.5 µM) of exogenous oxidizing agents were applied. At 24 hpi, IFA and IFU analyses were performed to quantify the effect of the oxidants on chlamydial growth in the absence of *ahpC* activity. Under conditions of *ahpC* knockdown, even sublethal concentrations (62.5 µM) of H_2_O_2_, CHP, or TBHP further reduced the inclusion size, and the IFU data revealed a decrease from >90% to <30% in comparison to the untreated control (Fig. 6A, 6B, and S5). Reduced expression of *ahpC* also affected the survival of *Chlamydia* against peroxynitrite. Notably, there was no significant change in the NT control strain in the uninduced and induced samples in response to oxidizing agents as assessed by IFU and IFA (Fig. 6C, 6D, and S5). These data further support that the chlamydial AhpC is a critical antioxidant enzyme in these bacteria.

**Fig. 6.**
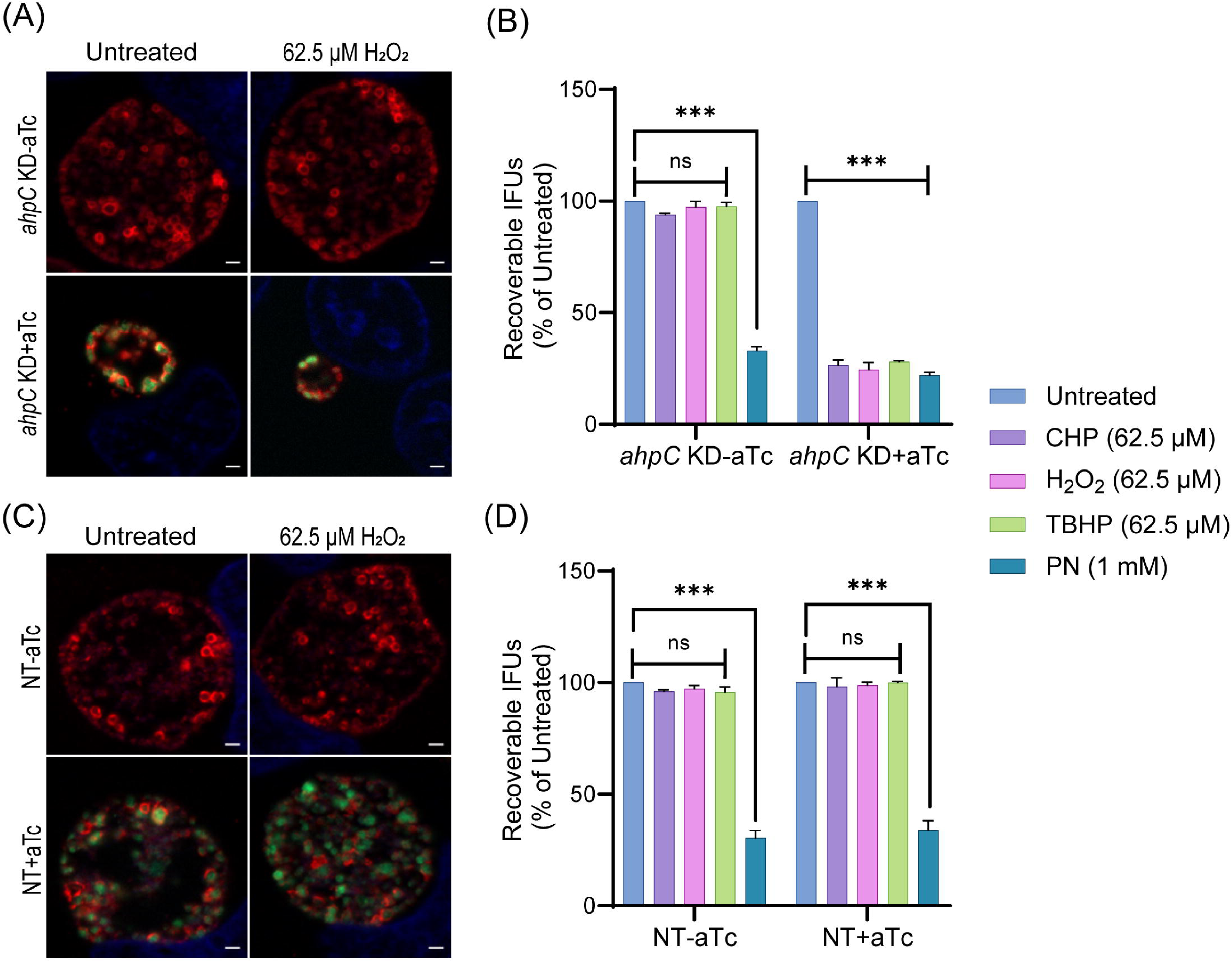
C*h*lamydia is hypersensitive to oxidizing agents in *ahpC* knockdown condition. IFA of *ahpC* KD (A) or NT (C) treated with 62.5 µM H_2_O_2_. dCas12 expression was induced or not at 10 hpi with 1 nM aTc, treated or not with H_2_O_2_ at 16 hpi for 30 min, and allowed to grow until 24 hpi. Coverslips were fixed with methanol at 24 hpi and stained major outer membrane protein (MOMP), Cpf1 (dCas12), and DAPI. Scale bars = 2 µm. Images were captured using a Zeiss Axio Imager Z.2 with Apotome2 at 100x magnification. Representative images from three biological replicates are shown. IFU analysis of *ahpC* KD (B) or NT (D) following treatment with oxidizing agents, CHP-Cumene hydroperoxide, H_2_O_2_-Hydrogen peroxide, TBHP-Tert-butyl hydroperoxide, or PN-Peroxynitrite. dCas12 expression was induced or not, and samples were treated or not as mentioned in the legend of Fig. 3B. IFUs of treated samples were calculated as percentage of respective untreated samples. \*\*\**p*< 0.0001 vs untreated sample by using two-way ANOVA. Data represent three biological replicates.

### Complementation restores the growth and resistance to low levels of peroxide stress of the *ahpC* knockdown strain

To validate that the impaired growth, altered inclusion morphology, and enhanced sensitivity to peroxides were due to the decreased level/activity of AhpC during knockdown, we generated a complemented strain to restore *ahpC* expression during knockdown. For the construction of the *ahpC* complementing plasmid, the *ahpC* gene was cloned and transcriptionally fused 3’ to the dCas12 in the pL12CRia(*ahpC*) knockdown plasmid. Here, the complementing *ahpC* allele is also under the control of the aTc induced P*tet* promoter and is co-expressed with the aTc inducible dCas12. Consequently, the *ahpC* knockdown effect is ablated, and the observed phenotypes should be restored. After inducing *dCas12-ahpC* expression with aTc, the resultant strain was verified by RT-qPCR. Increased transcripts for *ahpC* were quantified under these conditions, indicating successful complementation of the knockdown effect (Fig. 7A). Of note, the *ahpC* transcript levels remained elevated in comparison to the uninduced control at the 24 hpi time point. After confirming this strain, we measured genomic DNA to quantify total bacteria and performed IFA and IFU assays to examine if complementation restored the phenotypes observed during *ahpC* knockdown. These assays revealed normal inclusion morphology, gDNA levels, and EB progeny production in the complemented strain, indicating successful complementation of the knockdown phenotype (Fig. 6B, C, and D). Of note, a C-terminal 6xHis tagged AhpC was not capable of complementing the knockdown phenotype (data not shown), indicating a requirement for a free C-terminus in the function of AhpC in *Chlamydia*. Previous studies in other bacteria have revealed that the C-terminal residues in AhpC play a crucial role in the structural stability and enzymatic activity of AhpC (Dip et al., 2014; Feng et al., 2020; Wan et al., 2021).

**Fig. 7.**
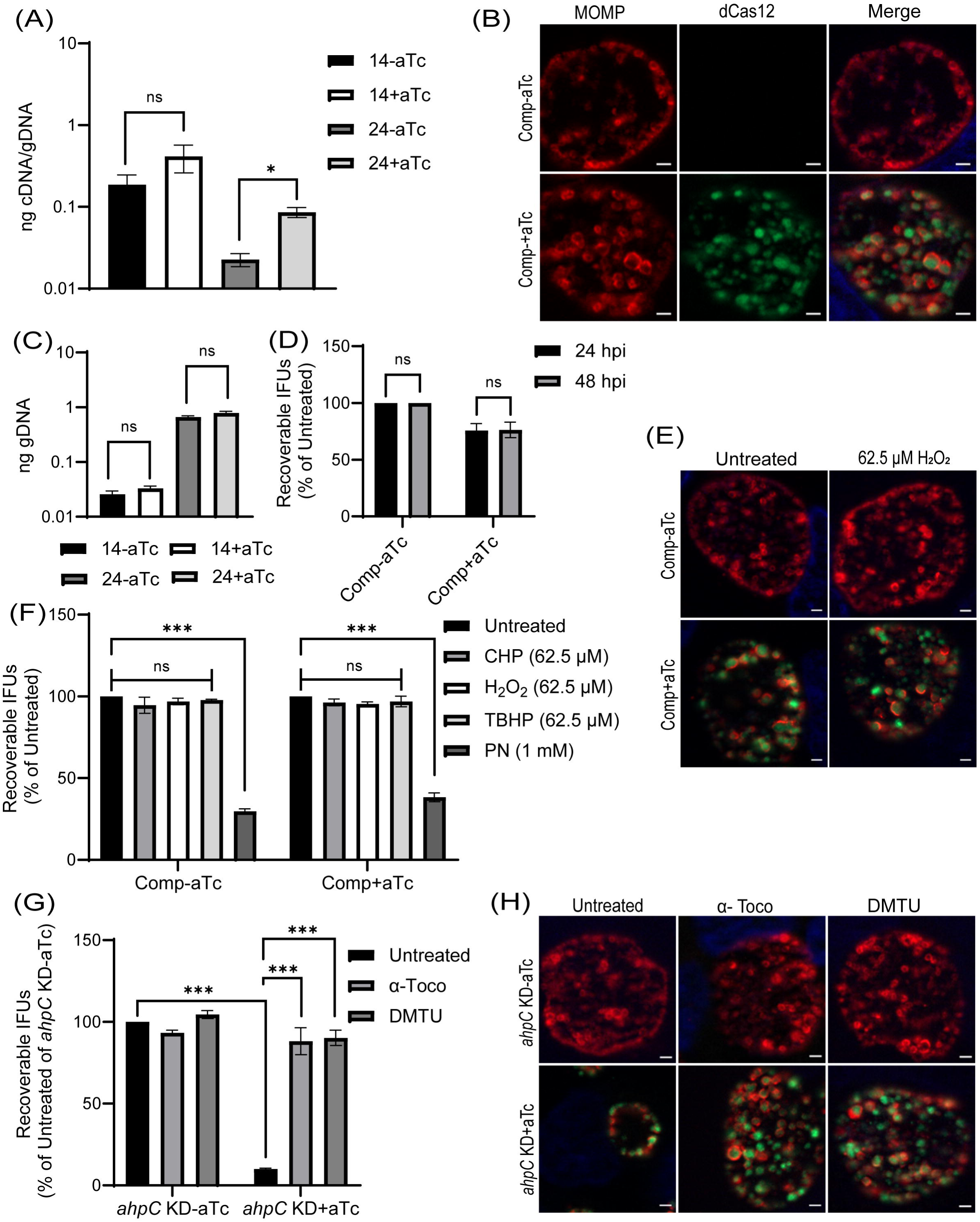
Complementation of the phenotypes observed in the *ahpC* knockdown. (A) Confirmation of complementation (comp) of *ahpC* knockdown by RT-qPCR. Samples were processed and quantified as mentioned previously in the legend of Fig. 4A. Values were plotted on a log scale. \**p*< 0.01 vs uninduced sample using ordinary one-way ANOVA. Data represent three biological replicates. (B) IFA of comp strain was performed at 24 hpi, and staining and imaging were performed as mentioned in the legend of Fig. 4B. Scale bars, 2 µm. Representative images from three biological replicates are shown. (C) Genomic DNA quantitation was performed by qPCR. Construct expression was induced or not at 10 hpi, and gDNA was harvested at 14 and 24 hpi and plotted on a log scale. Statistical analysis was calculated using ordinary one-way ANOVA. Data represent three biological replicates. (D) IFU analysis of comp strain. Statistical analysis was calculated using multiple paired t test. Data represent three biological replicates. (E) IFA of comp strain following treatment with 62.5 µM H_2_O_2_. Samples were treated, stained, and images acquired as mentioned in the legend of Fig. 6A. Scale bars = 2 µm. Representative images from three biological replicates are shown. (F) IFU analysis of comp strain following treatment with oxidizing agents. Experiments were performed as mentioned in the legend of Fig. 6B. IFUs were calculated as percentage of respective untreated samples. \*\*\**p*< 0.0001 vs untreated sample by using two-way ANOVA. Data represent three biological replicates. (G) *ahpC* knockdown growth defect rescued by ROS scavengers. IFU analysis of *ahpC* knockdown treated with or without scavengers, α-Tocopherol (100 µM) and DMTU (10 mM), as mentioned in materials and methods. IFUs were calculated as a percentage of the untreated, uninduced sample. \*\*\**p*< 0.0001 vs untreated, uninduced or induced sample by using two-way ANOVA. Data represent three biological replicates. (H) IFA of *ahpC* knockdown treated with or without scavengers. Experimental conditions were similar as in section (G). Staining and imaging were performed as mentioned in Fig. 6A. Representative images from three biological replicates are shown.

We next assessed whether complementation of the knockdown phenotype could also restore the resistance to low levels of peroxide stress. As in previous experiments, a sublethal concentration of oxidizing agents was added for 30’ at 16 hpi after having induced *dCas12-ahpC* expression at 10 hpi. Consistent with the growth parameters (Fig. 7B-D), the complemented strain showed wild-type responses in these conditions (Fig. 7E, 7F, and S5). These data indicate that the enhanced susceptibility of the *ahpC* knockdown strain to oxidizing agents was due to the reduced levels of *ahpC* and not an indirect effect of knockdown.

We predicted that the growth defects resulting from higher production of ROS during *ahpC* knockdown could be rescued by treating the *ahpC* KD with ROS scavengers. To test this prediction, we utilized two characterized ROS scavengers, DMTU (*N,N’-dimethylthiourea*) and α-tocopherol, which scavenge H_2_O_2_ and peroxyl radical, respectively (Hollander-Czytko et al., 2005; Kiffin et al., 2006; Walch et al., 2015). First, the effect of ROS scavengers on uninfected and infected HeLa cells was investigated using a viability assay. This assay revealed that 10 mM DMTU and 100 µM α-tocopherol had no adverse effects on uninfected or EV-infected HeLa cells (data not shown). These same concentrations were tested on WT *Ctr* L2 infected HeLa cells in untreated and 500 µM H_2_O_2_ treated conditions to test the potency of these scavengers to rescue growth defects associated with this concentration of oxidizing agent. Scavengers were added at 9.5 hpi and washed away at 16 hpi - the time of addition of H_2_O_2_ in the respective samples. At 16.5 hpi, after 3 wash steps, scavengers were added again with fresh media, and, at 24 hpi, bacterial growth and morphology were assessed using IFU and IFA assays. Neither scavenger had a significant impact on chlamydial morphology or infectious progeny. In the case of peroxide (500 µM H_2_O_2_) treated samples, scavengers restored IFUs from ∼50% to ∼100%, and inclusion size was also recovered (Fig. S6A and B).

Next, the effect of the ROS scavengers under *ahpC* knockdown conditions was assessed. In this experiment, scavengers were added at 9.5 hpi to provide the protective effect of scavengers before reducing the activity of *ahpC*. At 10 hpi, knockdown was induced or not with 1 nM aTc, and, at 14 hpi, scavengers were added again. At 24 hpi, IFU and IFA samples were collected to examine bacterial growth and morphology to assess the effect of ROS scavenging on the *ahpC* knockdown phenotype. As shown in Fig. 7G and 7H, both ROS scavengers had a positive impact on restoring growth of the *ahpC* knockdown strain; inclusions were larger, and IFUs were increased from <10% to >90% in the presence of scavengers in the induced samples. These data provide compelling evidence that the adverse effects of *ahpC* knockdown are due to increased ROS accumulation in this strain.

### Chlamydial developmental cycle progression is altered by *ahpC* knockdown/ overexpression

Whereas the overexpression of AhpC delayed overall developmental progression and was consistent with our hypothesis, the results with the *ahpC* knockdown strain appeared to refute our hypothesis given the apparent decrease in IFUs and smaller inclusion sizes after knockdown. To investigate the effects of changes in redox potential on chlamydial developmental cycle progression, we performed a transcriptional analysis of well-characterized late-cycle genes associated with secondary differentiation (*hctA*, *hctB*, *glgA*, *tsp,* and *omcB*) (Barry et al., 1993; Belland et al., 2003; Brickman et al., 1993; Gehre et al., 2016; Newhall, 1987; Swoboda et al., 2023). In the uninduced samples, transcript levels of all the tested late genes were higher at 24 hpi compared to 14 hpi, indicating their normal expression during the late stage of the developmental cycle. In comparison to the uninduced control, under conditions of *ahpC* knockdown, significantly higher expression of *hctA*, *hctB*, *omcB, tsp,* and *glgA* was observed at the mid-developmental cycle timepoint of 14 hpi (Fig. 8A). At 24 hpi, expression of these genes was comparable between the uninduced and induced conditions in the *ahpC* knockdown strain. We subsequently performed RNA sequencing from the uninduced and induced *ahpC* knockdown strain at 14 hpi. Experimental conditions were same as described for the RT-qPCR study, and samples were processed as mentioned in materials and methods. RNA-seq results were statistically analyzed by the UNMC Bioinformatics Core facility (Table S2). We further categorized these genes based on significant difference (p<0.05) and fold-change (≥1.5). Using these stringent parameters, we identified 161 up-regulated genes and 145 down-regulated genes in the *ahpC* knockdown conditions (Table S2). Here, we observed a global increase in late gene transcripts at this mid-developmental cycle timepoint (Fig. 8B and Table 1), indicating that increased oxidation results in earlier increases in late gene transcripts. In contrast, the complementation strain showed no increase in the expression of these tested late genes at 14 hpi, and *tsp* and *glgA* transcripts were reduced (Fig. S7A). Consistent with our model, the overexpression of *ahpC* resulted in significantly lower expression of *hctA*, *hctB*, *omcB, tsp*, and *glgA* at 14 and 24 hpi (Fig. S7B), suggesting a delayed transition to EBs.

**Fig. 8.**
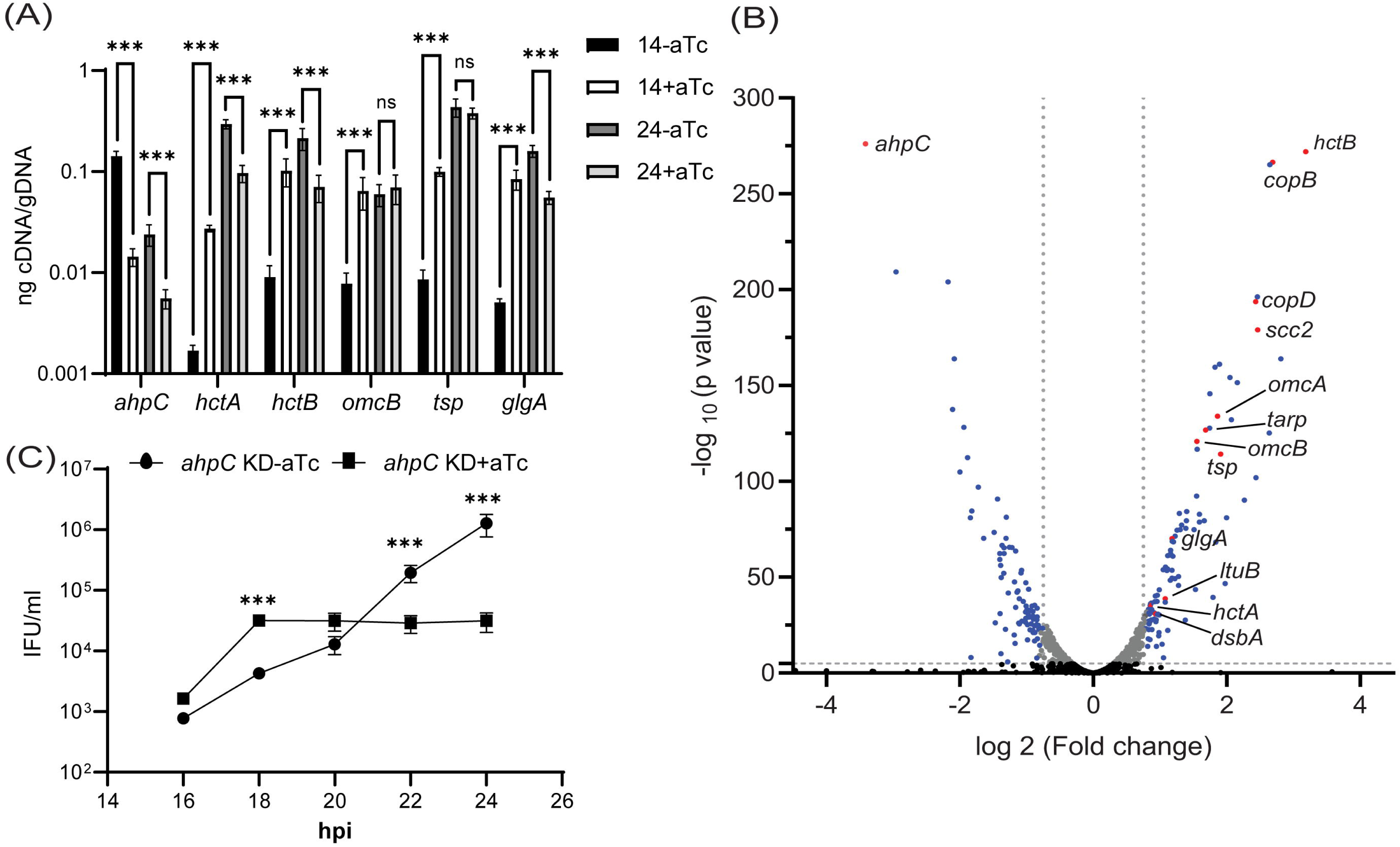
Effects of *ahpC* knockdown/overexpression on chlamydial developmental cycle progression. (A) RT-qPCR analysis of late cycle genes (*hctA*, *hctB*, *omcB*, *tsp*, and *glgA*) in *ahpC* knockdown. Experimental conditions were the same as mentioned in the legend of Fig. 4A. Quantified cDNA was normalized to gDNA, and values were plotted on a log scale. \*\*\**p*< 0.0001, \*\**p*< 0.001, \**p*< 0.01 vs uninduced sample by using two-way ANOVA. Data represent three biological replicates. (B) Volcano plot of RNA-sequencing of *ahpC* knockdown. Experimental conditions were the same as mentioned in the legend of Fig. 4A. The volcano plot was prepared using GraphPad Prism software. The vertical dashed lines indicate a fold change of 1.7 as compared to the respective uninduced control. The horizontal dashed line indicates an p value of 0.05. The black dots represent genes not significantly different and grey dots represent significantly altered but fold change lesser than 1.5. Blue and red spots represent statistically significant altered genes with more than 1.5-fold change in transcription levels between uninduced and induced samples. Red dots indicate ahpC and canonical late genes also shown in Table 1. (C) One-step growth curve of *ahpC* knockdown. Samples were induced or not with 1 nM aTc at 10 hpi and harvested at 16, 18, 20, 22, and 24 hpi. IFUs recovered are displayed as log10 values. \*\*\**p*< 0.0001 vs uninduced sample by using two-way ANOVA. Data represent three biological replicates.

**Table 1:**
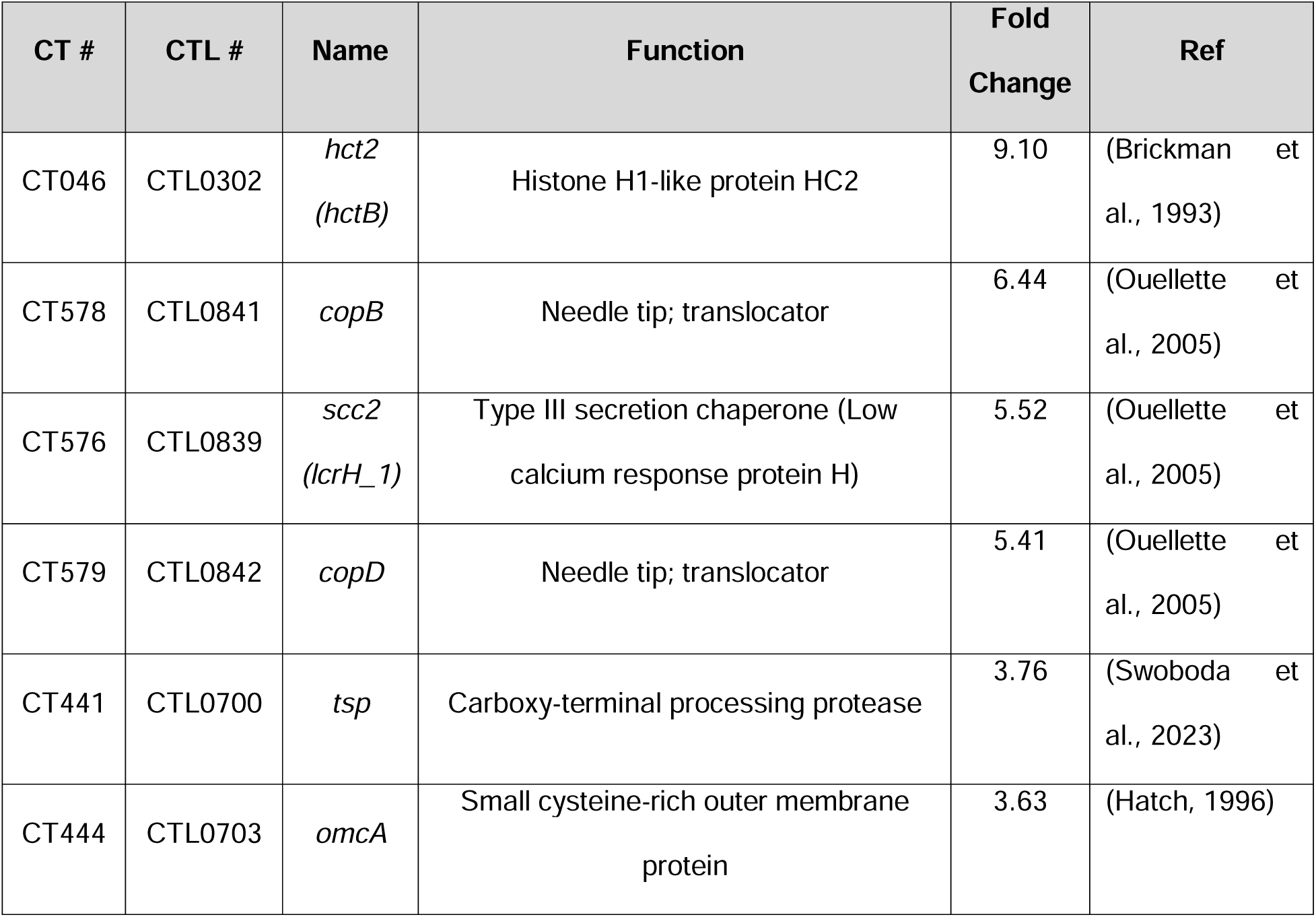
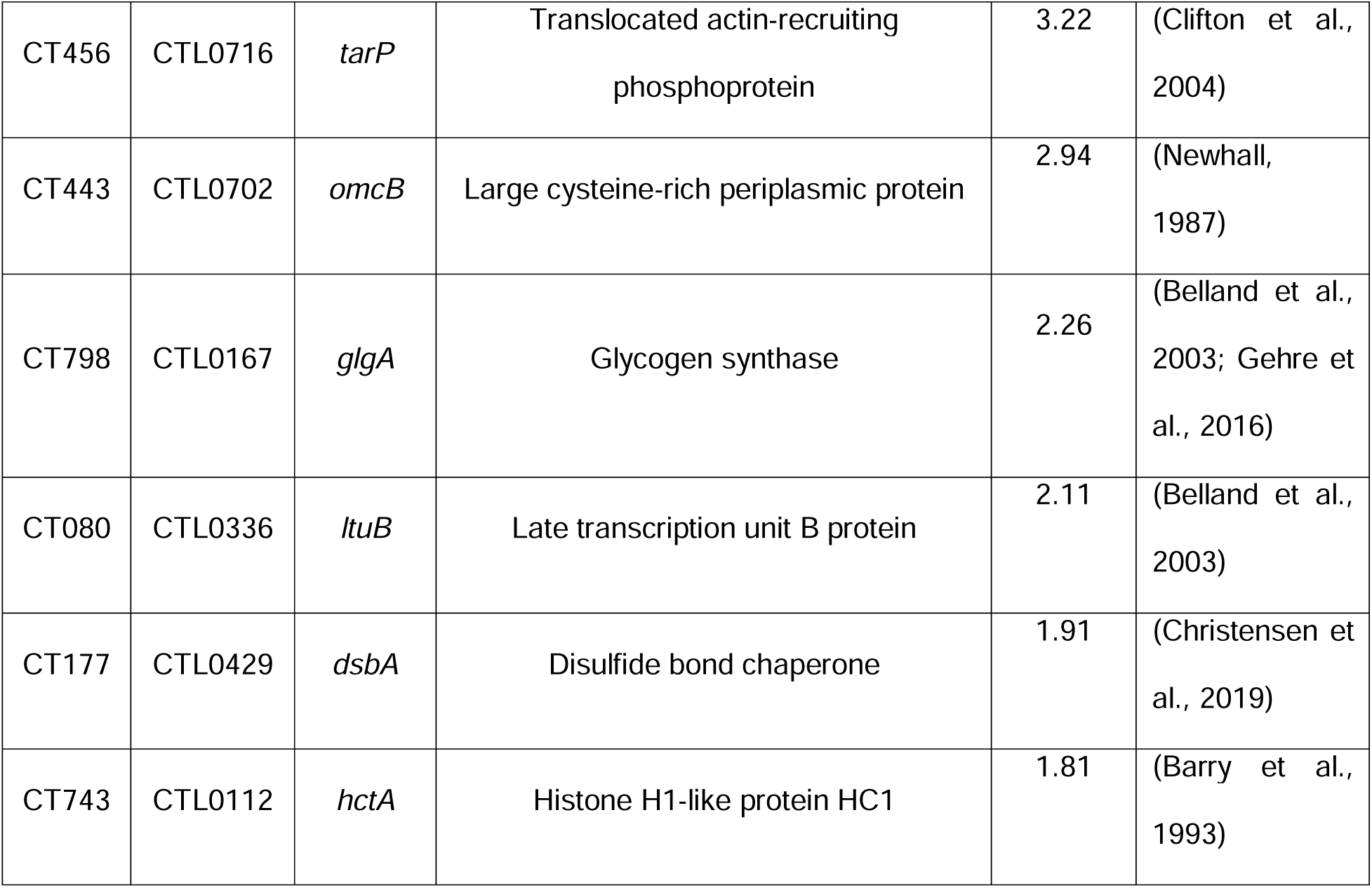
List of canonical late genes significantly increased during *ahpC* knockdown at 14 hpi.

Given the increase in late gene expression during *ahpC* knockdown at an earlier time point in the developmental cycle than normal, we reasoned that the increase in oxidizing conditions might prematurely trigger secondary differentiation and EB production. However, complicating such an analysis is that fewer RBs are present to convert to EBs, which would result in overall lower IFU yields as we had measured at 24 and 48 hpi (Fig. 4D). Nonetheless, to assess EB production more rigorously at earlier time points in the developmental cycle, we performed a growth curve analysis to precisely measure IFUs at two-hour intervals from 16 to 24 hpi. HeLa cells were infected, and knockdown was induced or not at 10 hpi with 1 nM aTc. At the indicated time points, IFU samples were collected and quantified. Consistent with our prediction, this experiment revealed higher IFUs (i.e., EBs) at 16 and 18 hpi during *ahpC* knockdown compared to the uninduced samples (Fig. 8C). This difference was statistically significant at 18 hpi. However, further EB production was stalled, with the uninduced strain continuing to produce EBs such that, by 24 hpi, there were significantly higher EB yields under these conditions (as noted in Fig. 4D). These data show that the phenotypic consequence of higher expression at 14 hpi of genes functionally related to EBs is the concomitant earlier production of EBs. These data underscore that reduced activity of *ahpC* causes earlier secondary differentiation in *C. trachomatis*.

### *ahpC* knockdown activates transcription of late-cycle genes when bacterial replication is blocked

We detected earlier expression of EB-related genes during *ahpC* knockdown and were curious if the *ahpC* knockdown condition could activate these genes under conditions when the chlamydial developmental cycle is blocked. Pathogenic *Chlamydia* species undergo a polarized cell division process, in which peptidoglycan is transiently synthesized only at the division septum (Abdelrahman et al., 2016; Cox et al., 2020; Ouellette et al., 2020). As a result, during penicillin treatment, division of RBs is blocked (Moulder, 1993; Moulder et al., 1956; Ouellette et al., 2020). However, the bacteria continue to grow in size resulting in aberrantly enlarged RBs (Barbour et al., 1982; Matsumoto & Manire, 1970; Ouellette et al., 2012), in which EB-related genes are not transcribed and production of EBs is inhibited (Ouellette et al., 2006; Panzetta et al., 2018).

To examine this, we generated an *ahpC* KD strain with spectinomycin resistance (*ahpC* KD-spec) and validated its phenotype as being the same as the penicillin-resistant *ahpC* KD strain. This new strain, *ahpC* KD-spec, was used to infect HeLa cells, and knockdown was induced or not at 10 hpi with 1 nM aTc. At the same time, samples were treated or not with 1 unit per mL of penicillin (Pen). RNA samples were harvested at 16 and 24 hpi and processed for RT-qPCR (Fig. S8B-D). IFA controls from these different conditions demonstrated the expected phenotypes (Fig. S8A). For example, Pen treatment caused aberrantly enlarged RBs, irrespective of *ahpC* KD, whereas *ahpC* KD itself caused smaller inclusions. Similarly, *ahpC* KD resulted in reduced *ahpC* transcripts in the induced conditions irrespective of the presence of Pen (Fig. S8B). Next, we quantified transcript levels for the late genes *hctA* and *hctB* in these different conditions (Fig. S8C and D). As expected, in the uninduced and untreated condition, both transcripts had higher expression at 24 hpi, indicating their regular developmental expression during the late stage of the developmental cycle. Consistent with prior observations (Ouellette et al., 2006), levels of these genes in uninduced + Pen conditions was low. As we previously noted (Fig. 8A), the transcripts of *hctA* and *hctB* are higher in the induced than uninduced samples at 16 hpi in the absence of Pen. Interestingly, in the induced + Pen condition, these genes had higher expression at 24 hpi than the uninduced + Pen control. At 16 hpi, their expression was either higher or similar between both samples (i.e., induced + Pen and uninduced + Pen). Collectively, these data support our observation that *ahpC* knockdown activates the transcription of late-cycle genes – even under conditions where developmental cycle progression is blocked.

## DISCUSSION

Many bacteria use redox sensing as a mechanism to control gene expression and subsequent morphologic transitions. Typically, these organisms use a redox-sensitive transcription factor, such as OxyR to sense these changes in redox conditions and alter gene expression appropriately. For example, in *Pseudomonas aeruginosa*, OxyR is involved in regulation of cellular metabolism, in addition to oxidative stress defense (Wei et al., 2012). In *Caulobacter crescentus*, redox has been reported as a regulator of metabolism and cell cycle progression (Hartl et al., 2020; Narayanan et al., 2015). Thus, the ability of bacteria to assess oxidative stress (imbalanced level of ROS) and to make physiological adjustments by altering gene expression patterns that favor their growth, and development is critical for their survival.

ROS generation is an inevitable condition for pathogens growing in aerobic conditions, and resistance against oxidative stress is a key survival mechanism (Fang, 2011). Hence, pathogens have evolved detoxifying proteins, such as peroxiredoxins, to eliminate ROS (Dip et al., 2014; Wang et al., 2004; Yang et al., 2002). AhpC has been reported to have a crucial role in bacterial physiology, survival, and virulence by scavenging ROS and RNI (Cosgrove et al., 2007; Kimura et al., 2012; Loprasert et al., 2003; Oh & Jeon, 2014). In the process of detoxification of peroxides and peroxynitrite, AhpC is converted into an oxidized dimer, requiring alkyl hydroperoxide reductase AhpF or AhpD to regenerate its activity (Koshkin et al., 2004; Poole & Ellis, 1996; Wong et al., 2017). *C. trachomatis* encodes AhpC (Ct603/CTL0866) but lacks any annotated homologs of AhpF or AhpD. Therefore, it remains an open question how AhpC activity is regulated. Some studies have mentioned *ahpC* as an iron-responsive gene in *Chlamydia* (Brinkworth et al., 2018; Pokorzynski et al., 2019), but there is no detailed investigation to date about the role of this crucial antioxidant in the growth and development of *Chlamydia.* Our study is the first to characterize a function of AhpC in chlamydial biology.

An essential aspect of the scavenging activity of AhpC for many bacterial pathogens in which it has been studied is that it works best with endogenous (i.e., low) levels of H_2_O_2_. These bacteria, such as *Staphylococcus aureus* and *Yersinia pseudotuberculosis*, encode catalases, which have high Km for hydrogen peroxide that serve a predominant role in scavenging exogenous H_2_O_2_ (Cosgrove et al., 2007; Wan et al., 2021). Catalases can detoxify H_2_O_2_ at high levels (millimolar levels) and are crucial in responding to external H_2_O_2_ stress (Mishra & Imlay, 2012). The *ahpC* mutants in these bacteria become sensitive to organic peroxides but resistant to H_2_O_2_ due to higher catalase activity as a compensatory response to the lack of AhpC (Antelmann et al., 1996; Mongkolsuk et al., 2000; Ochsner et al., 2000). Further, simultaneous mutations in both catalase and *ahpC* displayed drastically enhanced sensitivity to all oxidizing agents tested (Cosgrove et al., 2007; Ezraty et al., 2017; Seaver & Imlay, 2001). *C. trachomatis* does not encode a catalase gene (Boncompain et al., 2014; Rusconi & Greub, 2013), and hypersensitivity of *ahpC* knockdown to both inorganic and organic peroxides indicates the absence of catalase and establishes that AhpC is the primary scavenger of ROS in *C. trachomatis*. Contrary to previous studies, we did not observe significant effects of PN with altered AhpC expression. One possible explanation may be our study’s concentration (1 mM) of PN. Higher concentrations of PN could not be used due to toxic effects on the host cell (data not shown). The other possibility may be the presence of some other unknown mechanism(s) to detoxify PN. For example, *Chlamydia* also encodes a superoxide dismutase (SOD) that may prevent the accumulation of the necessary precursors that are needed for PN production. We are currently investigating the function of the chlamydial SOD enzyme.

Resistance to different oxidants as a result of overexpression of *ahpC* (Fig. 3 and S3) is consistent with studies in other bacteria (Sherman et al., 1996; Zuo et al., 2014), signifying that AhpC is the principal defense mechanism against oxidative stress in *Chlamydia*. During *ahpC* knockdown, the attenuated activity of AhpC severely affected the growth of *Chlamydia* (Fig. 4). Notably, the observed morphology of inclusions during *ahpC* knockdown is strikingly similar to *Chlamydia* treated with a high concentration (1 mM) of oxidizing agents used (Fig. 4 and Fig S2), suggesting both conditions result in increased oxidative stress to the organism. This was further supported by the increased sensitivity of *ahpC* KD to sublethal concentrations of oxidants (Fig. 6 and S4), consistent with previous data from other bacteria (Cosgrove et al., 2007; Seaver & Imlay, 2001; Storz et al., 1989; Zhang et al., 2019). In *ahpC* KD, the ROS level was dramatically higher (Fig. 5 and S4), similar to previous studies in other bacterial systems (Zhang et al., 2019). Further, the addition of ROS scavengers to the culture medium rescued the negative phenotypes associated with *ahpC* knockdown (Fig. 7G and H). This strongly suggests that, in the absence of AhpC, higher amounts of reactive oxygen species accumulate in *Chlamydia* and that the negative impact on growth during *ahpC* knockdown is due to highly oxidized conditions in the organism. Moreover, for the complete restoration of inclusion size, multiple doses of scavengers were required throughout the experiment, further emphasizing that the internal chlamydial environment is oxidized and/or susceptible to oxidation in the absence of AhpC. These findings suggest that AhpC in *Chlamydia* is indispensable, having a functional role even more extensive than previously reported in other bacteria.

In further support of this observation, we noticed upregulated transcripts of late-cycle genes such as *hctA*, *hctB*, *glgA, omcA, omcB*, *dsbA*, and *tsp* at an earlier time (14 hpi) in the chlamydial developmental cycle as a result of reduced AhpC activity (Fig. 8). Notably, in the developmental cycle of *C*. *trachomatis*, only RBs are present at 14 hpi, and secondary differentiation starts after 16 hpi (Abdelrahman & Belland, 2005). Expression of most of the late-cycle genes occurs after this time in the developmental cycle (Belland et al., 2003). These late-cycle genes are well characterized for their association with EBs. The *hctA* and *hctB* genes encode histone-like proteins responsible for chromosomal condensation during the differentiation of RBs into EBs. The *hctA* gene is among the first to be transcribed in the late stage (Chiarelli et al., 2020). Tsp is a periplasmic protease thought to be crucial in the degradation of RB-specific periplasmic proteins during secondary differentiation in *C. trachomatis* (Swoboda et al., 2023). OmcA, OmcB, and DsbA are responsible for the rigid cell wall and osmotic stability of the EBs (Christensen et al., 2019; Hatch, 1996; Newhall, 1987). GlgA is key enzyme for glycogen accumulation in the inclusion lumen at late stages of the developmental cycle (Gehre et al., 2016). One crucial point to consider is the redox status of *Chlamydia*, 14 hpi is the time when reduced RBs predominate with virtually no oxidized EBs detectable. However, we established that *ahpC* knockdown shifts the redox status of the bacteria towards oxidation. Hence, the earlier and significantly higher detection of transcripts for these late-cycle genes indicate earlier secondary differentiation in the *ahpC* knockdown. Importantly, *ahpC* complementation during knockdown restored the regular developmental expression of these late-cycle genes, further supporting that this phenotype was due to the highly oxidized conditions created by reduced AhpC activity. The conclusion of these data is that AhpC levels, and presumably activity, have a direct effect on secondary differentiation.

Secondary differentiation (differentiation from RBs to EBs) is an essential step for chlamydial growth and survival, but there is a dearth of information regarding the mechanisms of its regulation. EBs and RBs have significantly different proteomic repertoires (Saka et al., 2011; Skipp et al., 2005; Skipp et al., 2016), and our group has identified critical functions for the cytoplasmic ClpXP and ClpCP and periplasmic Tsp proteases during secondary differentiation (Pan et al., 2023; Swoboda et al., 2023; Wood et al., 2022).The *tsp* gene, in addition to another late-cycle gene, *hctB*, is transcriptionally regulated by sigma factor 28 (^28^) (Douglas & Hatch, 2000; Hatch & Ouellette, 2023). Another sigma factor, ^54^ has been linked to regulation of outer membrane components, type III secretion system components, and other genes typically expressed late in development (Hatch & Ouellette, 2023; Soules et al., 2020). In addition to the two minor sigma factors, ^28^ and ^54^, *Chlamydia* encodes one major sigma factor, ^66^, which is transcriptionally regulated by the relative levels of RsbV1 (antagonist) and RsbW (anti-sigma factor) (Thompson et al., 2015). This Rsb system senses ATP availability and has been proposed to regulate ^66^ (Thompson et al., 2015). A related study from Soules et al. (Soules et al., 2020) showed that TCA intermediates act as ligands for RsbU, thereby linking the Rsb system to the TCA cycle and ATP synthesis. Collectively, these data show that post-translational mechanisms drive secondary differentiation in *Chlamydia*. However, no definitive “switch” that triggers this step has been identified.

How might increasing oxidation regulate secondary differentiation? *Chlamydia* lacks an identifiable ortholog of OxyR so some other mechanism must be employed. We speculate that increasing ROS production from the metabolic activity of the RB will overcome the innate and stochastic activity of AhpC to detoxify ROS. This could either directly alter the activity of specific proteins or lead to the accumulation of damaged proteins that in turn triggers secondary differentiation. These possibilities are not mutually exclusive. We propose a simple model to explain this (Fig. 9). As secondary differentiation is asynchronous and RBs divide through an asymmetric budding mechanism (Ouellette et al., 2020), AhpC as well as proteins impacted by ROS levels are unevenly distributed between mother and daughter cell. This difference will lead some RBs to breach an oxidative threshold sooner, allowing activation of late genes and secondary differentiation earlier than other RBs. Consistent with this, the higher expression of genes functionally related to EBs and EB production during *ahpC* KD at an early stage of the chlamydial developmental cycle indicates earlier secondary differentiation as an outcome of diminished activity of *ahpC* in *C. trachomatis*. Consequently, increased late gene transcription should have the phenotypic effect of causing earlier production of EBs. Indeed, we quantified more IFUs in the *ahpC* knockdown at 16 and 18 hpi compared to the uninduced control condition. During *ahpC* KD, higher amounts of ROS accumulation in the bacterium create highly oxidized conditions. In contrast, *ahpC* overexpression scavenges ROS at a greater level leading to a more reducing environment. For the AhpC overexpression strain, the expression of late-cycle genes in the induced conditions compared to the uninduced control at 24 hpi is significantly lower with reduced production of EBs. These data support our hypothesis that the developmental cycle is delayed in these conditions with a concomitant delay in achieving the oxidative threshold, thus allowing RBs to continue to divide before committing to secondary differentiation. Taken together, our data directly link oxidation state and secondary differentiation in *Chlamydia*. Ongoing studies are focused on characterizing redox-sensitive chlamydial proteins to understand which specific factors drive the shift from RBs to EBs (or EBs to RBs) in *C. trachomatis*.

**Fig. 9.**
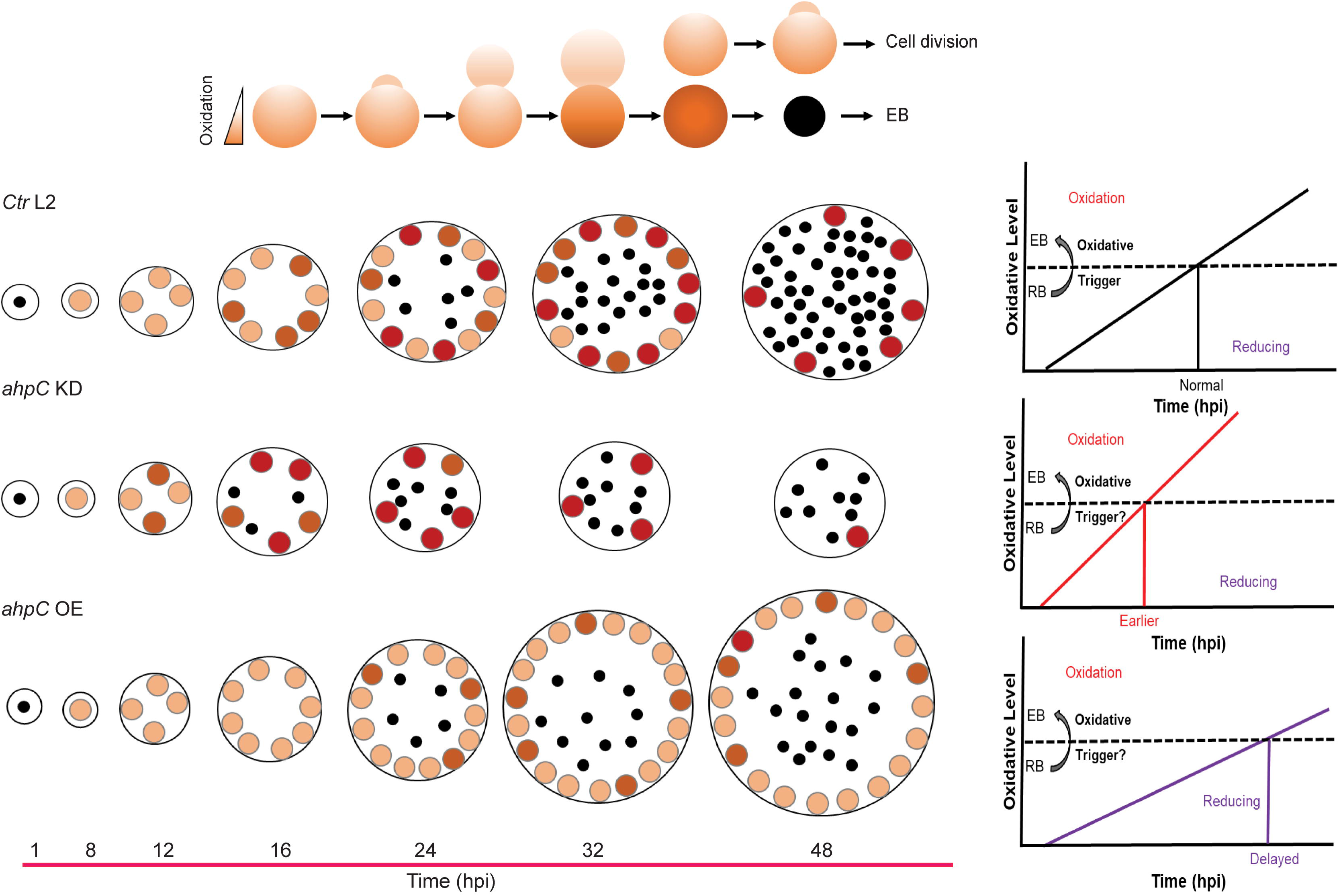
Altering the activity of AhpC in *Ctr* L2 impacts its developmental cycle progression. (Top) In *Chlamydia*, secondary differentiation is asynchronous and RBs divide through an asymmetric budding mechanism. In such conditions, either the mother or daughter cell may inherit more oxidized proteins (represented by a darker shade), which can then impact whether a given RB will divide again or undergo secondary differentiation. (Bottom) The black dots represent EBs, the orange circles show RBs. The developmental cycle of wild-type *C. trachomatis* (*Ctr* L2) is shown. In *ahpC* KD, highly oxidized conditions lead some RBs to cross the oxidative threshold sooner, allowing activation of late genes and secondary differentiation earlier than other RBs. In *ahpC* overexpression, a reducing environment results in a delay in achieving the oxidative threshold, thus allowing RBs to continue to divide before committing to secondary differentiation.

## MATERIALS AND METHODS

### Strains and cell culture

For chlamydial transformation, McCoy mouse fibroblast cells (kind gift of Dr. Harlan Caldwell (NIH/NIAID)) were used. Human cervix adenocarcinoma epithelial HeLa cells (kind gift of Dr. Harlan Caldwell (NIH/NIAID)) were used for RT-qPCR, immunofluorescence assays (IFA), inclusion forming unit assays (IFU), viability assays, oxidative stress, RNA-seq, ROS measurements, and ROS scavenger assays. Both cell types were routinely grown and passaged in Dulbecco’s modified Eagle’s medium (DMEM; Gibco, Waltham, MA) supplemented with 10% fetal bovine serum (FBS; Sigma, St. Louis, MO) and 10 µg/mL gentamicin (Gibco, Waltham, MA) at 37°C and 5% CO2. All strains were verified to be Mycoplasma-negative using LookOut mycoplasma PCR detection kit (Sigma). For chlamydial transformations, *Chlamydia trachomatis* serovar L2 EBs lacking the endogenous pL2 plasmid (kind gift of Dr. Ian Clarke, University of Southampton) were used (Wang et al., 2011). Wild-type, density gradient-purified *Chlamydia trachomatis* 434/Bu (ATCC VR902B) EBs were used for sensitivity to oxidizing agents and ROS scavenger assays. Molecular biology reagents, oxidizing agents, CellROX Deep Red dye, and scavengers were purchased from Thermo Fisher unless otherwise noted.

### Plasmid construction

The primers, gBlock gene fragments, plasmids, and bacterial strains used for molecular cloning are listed in Table S1 in the supplemental material. Constructs for chlamydial transformation were cloned using high-fidelity (HiFi) cloning system from New England BioLabs (NEB). Primers were designed using the NEBuilder online primer generation tool (https://nebuilderv1.neb.com). For the overexpression strain, the *ahpC* gene was amplified by PCR with Phusion DNA polymerase (NEB) using *C. trachomatis* serovar L2 434/Bu genomic DNA as a template. The PCR product was purified using a PCR purification kit (Qiagen, Hilden, Germany). The HiFi assembly reaction was performed as per the manufacturer’s instructions in conjunction with the pBOMBDC plasmid digested with Fast Digest EagI and KpnI enzymes and dephosphorylated with FastAP (Thermo Fisher, Waltham, MA). The HiFi reaction mix was transformed into *E. coli* 10-beta (NEB). Plasmids were first confirmed by restriction enzyme digestion, and final verification of insert was performed using Sanger sequencing. A dCas12-based CRISPRi approach was used for *ahpC* knockdown generation (Ouellette et al., 2021). The pBOMBL12CRia plasmid having anhydrotetracycline (aTc) inducible catalytically dead dCas12 protein was used as vector. A crRNA targeting the 5’ intergenic region of *ahpC* was designed and ordered as a presynthesized DNA fragment. For *ahpC* knockdown construct, 2 ng of the gBlock (Integrated DNA Technologies [IDT], Coralville, IA) listed in Table S1 was combined with 25 ng of BamHI-digested, alkaline phosphatase-treated pBOMBL12CRia(e.v.)::L2 in a HiFi reaction according to the manufacturer’s instructions (NEB). The plasmid was transformed into NEB 10-beta cells and verified by Sanger sequencing prior to transformation into *Chlamydia trachomatis*. To generate the complementation strain, the *ahpC* gene was amplified using primers listed in Table S1 and fused with the *ahpC* knockdown construct (i.e., pBOMBL12CRia (*ahpC*)) digested with the Fast Digest restriction enzyme SalI (Thermo Fisher) and alkaline phosphatase-treated, using the HiFi reaction as mentioned above.

### Chlamydial transformation

Chlamydial transformations were performed using a protocol described previously, with some modifications (Mueller et al., 2017). One day before transformation, 1 x 10^6^ McCoy cells were seeded in one 6-well plate, and two wells were used per plasmid transformation. Briefly, for each well of the 6-well plate, 2 µg of sequenced verified plasmids were incubated with 2.5 x 10^6^ *C. trachomatis* serovar L2 without plasmid (-pL2) EBs in 50 µL Tris-CaCl_2_ (10 mM Tris, 50 mM CaCl2, pH 7.4) at room temperature for 30 min. McCoy cells were washed with 2 mL Hank’s Balanced Salt Solution (HBSS; Gibco), and 1 mL HBSS was added back into each well. 1 mL of HBSS was added to each transformant mixture, and one well of a 6-well plate was infected using this transformation solution. Cells were centrifuged at 400 × g for 15 min at room temperature followed by 15 min incubation at 37°C. HBSS was aspirated and replaced with antibiotic-free DMEM. At 8 hpi, 1 µg/mL of cycloheximide and 1 or 2 U/mL of penicillin G or 500 μg/mL spectinomycin were added to the culture media. The infection was passaged every 48 h until a population of penicillin, or spectinomycin resistant, green fluorescent protein (GFP) positive *C. trachomatis* was established. The chlamydial transformants were then serially diluted to isolate clonal populations. These isolated populations were further expanded and frozen at - 80°C in a sucrose phosphate solution (2SP). To verify plasmid sequences, DNA was harvested from infected cultures using the DNeasy kit (Qiagen) and transformed into NEB 10-beta for plasmid propagation. Isolated plasmids were then verified by restriction digest and Sanger sequencing.

### Inclusion forming unit assay

Inclusion forming unit assay was performed to determine the infectious progeny (number of EBs) from a primary infection based on inclusions formed in a secondary infection. *C. trachomatis* transformants were infected into HeLa cells and induced or not with 1 nM aTc at 10 hpi. At 24 and 48 hpi, samples were harvested by scraping three wells of a 24-well plate in 2 sucrose-phosphate (2SP) solution and lysed via a single freeze-thaw cycle, serially diluted, and used to infect a fresh HeLa cell monolayer and allowed to grow for 24 h. Samples were fixed with methanol, stained with a goat antibody specific to *C. trachomatis* major outer membrane protein (MOMP; Meridian Biosciences, Memphis TN) followed by staining with donkey antigoat Alexa Fluor 594-conjugated secondary antibody (Invitrogen) and titers were enumerated using a 20x lens objective. All experiments were performed three times for three biological replicates. Induced values were expressed as a percentage of the uninduced values, which was considered as 100%.

### Immunofluorescence assay

HeLa cells were cultured on glass coverslips in 24-well tissue cultures plates at 2 x 10^5^ cells/well, infected with the relevant strains, and, at 10 hpi, samples were induced or not with 1 nM aTc. At 24 hpi, samples were fixed and permeabilized using 100% methanol. Organisms were stained with anti-MOMP (Meridian Biosciences) primary antibody for all the strains and primary mouse anti-Cpf1 (dCas12) (Sigma-Millipore) for knockdown or complementation samples. Donkey anti-goat Alexa Fluor 594-conjugated secondary antibody (Invitrogen) was used to visualize *Chlamydia* in all the samples, and donkey anti-mouse Alexa Fluor 488-conjugated secondary antibody (Invitrogen) was used for dCas12 expression. DAPI (Invitrogen) was used for visualization of host and bacterial cell DNA. These stained coverslips were mounted on glass slides using ProLong glass antifade mounting media (Invitrogen) and imaged using a 100x lens objective on a Zeiss Axioimager Z.2 equipped with Apotome.2 optical sectioning hardware and X-Cite Series 120PC illumination lamp using a 2MP Axiocam 506 monochrome camera.

### Nucleic acid extraction and RT-qPCR

HeLa cells were seeded in 6-well tissue culture plates, infected with the *C. trachomatis* transformants, and induced or not at 10 hpi with 1 nM aTc. For each condition, triplicate wells were used for simultaneous harvest of RNA (for transcript analysis by RT-qPCR), gDNA (to normalize RT-qPCR data and quantification of gDNA), and IFA (to verify the morphological changes in samples in the tested conditions). For RNA extraction, cells were rinsed with DPBS twice and lysed with 1 mL TRIzol (Invitrogen/Thermo Fisher) per well as per manufacturer’s instructions. 200 μL of chloroform was added to extract the aqueous layer containing total RNA, which was precipitated with isopropanol. A total of 10 µg of purified RNA was treated with TURBO DNase (Invitrogen/Thermo Fisher) according to the manufacturer’s instructions to remove DNA contamination. DNA-free RNA was used for cDNA synthesis using random nonamers (N9; NEB) and SuperScript III reverse transcriptase (Invitrogen/Thermo Fisher) following the manufacturer’s instructions. cDNA samples were diluted 10-fold with molecular biology-grade water and stored at -80°C. A total of 2.5 µL of each diluted cDNA sample was used per well of a 96-well qPCR plate. For each of three biological replicates, each sample was analyzed in triplicate on a QuantStudio 3 system (Applied Biosystems/Thermo Fisher) using the standard amplification cycle with melting curve analysis. For gDNA, one well of a 6-well plate per condition was scraped in 500 μL DPBS, split in half (i.e., 250 μL), and frozen at -80°C. Each sample was then thawed and frozen twice more for a total of three freeze/thaw cycles, and gDNA was extracted using the DNeasy DNA extraction kit (Qiagen) according to the manufacturer’s guidelines. The isolated gDNA was quantified and diluted down to 5 ng/mL prior to use in quantitative PCR (qPCR). A total of 2.5 µL of each diluted gDNA sample was mixed with 10 µL of PowerUp SYBR green master mix in a 96-well qPCR plate and was analyzed on a QuantStudio 3 system (Applied Biosystems). Each sample from each biological replicate was tested in triplicate. For each primer set used, a standard curve of gDNA was generated against purified *C. trachomatis* L2 genomic DNA, and the cDNA levels were normalized to gDNA levels or 16S rRNA for analysis. All experiments were performed three times for three biological replicates. At the same time, morphological differences were monitored in the IFA control, with samples fixed with methanol at the time of harvesting of RNA/gDNA. Staining and imaging were performed as described above.

### Enrichment, library preparation, and statistical analyses of RNA-sequencing samples

Three biological replicates were prepared and processed for RNA isolation as mentioned above. Enrichment of microbial RNA was done using MICROB*Enrich* and MICROB*Express* (Invitrogen/ThermoFisher) kits as per the manufacturer’s instructions. Samples were aliquoted, stored at -80°C and submitted to the UNMC Genomics Core. Before library preparation, quality control (QC) was done using Nanodrop and Fragment Analyzer (Advanced Analytical). Final libraries were quantified using Qubit DS DNA HS Assay reagents in Qubit Fluro meter (Life Technologies), and the size of the libraries was measured via Fragment Analyzer. Individual RNA-seq libraries were prepared using 400 ng of total RNA by Illumina Ribo-Zero plus Microbiome (Illumina, Inc. San Diego, CA) and were multiplexed and subjected to 100 bp paired read sequencing to generate >20 million pairs of reads per sample using Midoutput NextSeq 300 cycle kit by the UNMC Genomics Core facility (Table S2). The original fastq format reads were trimmed by fqtrim tool (https://ccb.jhu.edu/software/fqtrim) to remove adapters, terminal unknown bases (Ns), and low quality 3’ regions (Phred score < 30). The trimmed fastq files were processed by FastQC (Andrews, 2010). *Chlamydia trachomatis* 434/Bu bacterial reference genome and annotation files were downloaded from Ensembl (http://bacteria.ensembl.org/Chlamydia_trachomatis_434_bu/Info/Index). Sequencing data were analyzed by the Bioinformatics and Systems Biology Core (BSBC). The false discovery rate (FDR) and Bonferroni adjusted p values were provided to adjust for multiple-testing problem. Fold changes were calculated from the GLM, which corrects for differences in library size between the samples and the effects of confounding factors. The trimmed fastq files were mapped to *Chlamydia trachomatis* 434/Bu by CLC Genomics Workbench 23 for RNA-seq analyses.

### Data availability

The raw and processed RNA sequencing reads in fastq format have been deposited in Gene Expression Omnibus (GEO: www.ncbi.nlm.nih.gov/geo/) under Accession number GSE278846.

### Viability assay

Cell viability assays were performed using PrestoBlue (Invitrogen/Thermo Fisher). HeLa cells were seeded in 96 well plates and were infected or not with the indicated chlamydial strains (e.g., pBOMBDC.ev). Samples were treated or not as per the experimental parameters (e.g., different concentrations of oxidizing agents or scavengers). Medium without cells was used as the blank control. At the end point, 10% PrestoBlue (v/v) was added in the wells and incubated at 37°C for 30 min while protecting the plate from light. Fluorescence was read at excitation 560 nm and emission 590 nm using an Infinite M200 Pro (Tecan). The treated values were expressed as a percentage of the untreated values, which was considered as 100%. All experiments were performed three times for three biological replicates.

### Oxidizing agents’ susceptibility testing

Susceptibility of host cells and chlamydiae to different oxidizing agents was tested in HeLa cells infected with density gradient-purified *Chlamydia trachomatis* 434/Bu EBs or *C. trachomatis* transformants. For *C. trachomatis* transformants, expression of the construct was induced or not with 1 nM aTc at 10 hpi. At 16 hpi, different concentrations of oxidizing agents were added, and samples were incubated at 37°C for 30 min. These samples were washed three times with HBSS, fresh DMEM media was added in the wells, and cultures were allowed to grow until 24 hpi. At 24 hpi, inclusion forming unit assays and immunofluorescence analysis were performed as described above.

### ROS scavenging by chemical compounds

The scavenging capacity of different ROS scavengers was tested using HeLa cells infected with *Chlamydia trachomatis* 434/Bu EBs or the *ahpC* KD strain. To examine the protective effect of scavengers in the absence or presence of oxidative stress, *Chlamydia trachomatis* 434/Bu EBs infected HeLa cells were used. In these samples, scavengers were added or not at 10 hpi in respective wells. At 16 hpi, wells were washed three times with HBSS, and fresh DMEM containing 500 μM H_2_O_2_ or not was added. After 30 min, wells were washed three times with HBSS and fresh DMEM containing scavengers or not was added back in the respective wells, which were incubated until 24 hpi prior to collecting samples for IFU assay and IFA analysis. The effect of these scavengers was further tested in *ahpC* KD. HeLa cells were infected with *ahpC* KD, scavengers were added or not at 9.5 hpi before induction at 10 hpi with 1 nM aTc. At 14 hpi, scavengers were added again in the respective wells, and, at 24 hpi, samples were collected for IFU assay and IFA.

### ROS detection by CellROX Deep Red

Intracellular ROS levels were measured in uninfected and *ahpC* KD-infected HeLa cells using CellROX Deep Red dye (Invitrogen/Thermo Fisher). Samples were induced or not at 10 hpi with 1 nM aTc. Medium without cells was used as the blank control. At the end point, media was removed, samples were washed three times with DPBS and incubated with CellROX Deep Red dye for 30 min in the dark at 37°C. Fluorescence was read at excitation 644 nm and emission 665 nm using an Infinite M200 Pro (Tecan). Microscopic images were captured using live cells at 24 or 40 hpi. Experimental conditions were same as mentioned above. All experiments were performed three times for three biological replicates.

## ACKNOWLEDGMENTS

We thank Dr. H. Caldwell (NIH/NIAID) for providing eukaryotic cell lines and Dr. I. Clarke (University of Southampton) for providing the plasmidless strain of *C. trachomatis* serovar L2. We thank Dr. Elizabeth A. Rucks for critical feedback. We thank the members of the Rucks/Ouellette research group for thoughtful discussion of the material presented. Funding for this work was provided by the National Institutes of Health (NIH/NIAID) grants R01AI170688 and R21AI178150 to SPO. Authors thank the UNMC Genomics Core Facility and Bioinformatics and Systems Biology Core (BSBC) at UNMC for RNA-seq services. The UNMC Genomics Core Facility receives partial support from the National Institute for General Medical Science (NIGMS) INBRE - P20GM103427-19, as well as the National Cancer Institute (NCI) and The Fred & Pamela Buffett Cancer Center Support Grant-P30CA036727. The BSBC receives support from Nebraska Research Initiative (NRI) and NIH (2P20GM103427, 5P30CA036727, and 2U54GM115458).

## Supplementary Information

**Table S1** List of plasmids, strains, and primers used in this study.

**Table S2 List of differentially expressed genes in *ahpC* knockdown RNA-seq**

**Fig. S1** Viability assay of uninfected or infected HeLa cells treated with oxidizing agents H_2_O_2_ (A), CHP (B), TBHP (C), and PN (D). HeLa cells were infected or not with empty vector control, pBOMBDC.ev (EV), and treated or not with different concentrations of oxidizing agents at 16 hpi for 30 min. At 24 hpi, an end point viability assay was performed using PrestoBlue as mentioned in materials and methods. The treated values were expressed as a percentage of the untreated values, which were considered as 100%. Data represent three biological replicates.

**Fig. S2** Response of *C. trachomatis* L2 against oxidizing agents. (A) IFA of wild-type *Ctr* L2 exposed to oxidizing agents. Oxidizing agents’ treatment, staining, and imaging were performed as mentioned in materials and methods. Representative images from three biological replicates are shown. (B) IFU analysis of *Ctr* L2 post-exposure with oxidizing agents. Conditions were same as in section (A), and IFUs were harvested at 24 hpi. IFUs were calculated as percentage of untreated samples. \*\*\**p*< 0.0001 vs untreated sample by using one way ANOVA. Data represent three biological replicates.

**Fig. S3** Overexpression of *ahpC* provides resistance to peroxides in *Chlamydia.* IFA of *ahpC* (A) or EV (B) exposed to oxidizing agents (CHP, TBHP, and PN). Experiments were performed as mentioned in the legend of Fig. 3A. Representative images from three biological replicates are shown.

**Fig. S4** Intracellular ROS levels in *ahpC* knockdown at 40 hpi. HeLa cells were infected or not with *ahpC* knockdown, and construct was induced or not at 10 hpi with 1 nM aTc. At 40 hpi, samples were washed with DPBS and incubated with CellROX Deep red dye for 30 min in dark. Microscopic images of live cells on Zeiss Axio Imager Z.2 with Apotome2 at 100x magnification were captured. Scale Bars=10 µm. Representative images of three biological replicates are shown.

**Fig. S5** *Chlamydia* is hypersensitive to oxidizing agents as a result of reduced levels of *ahpC*. IFA of *ahpC* KD (A), NT (B), comp (C) treated with CHP, TBHP, and PN. Experimental conditions were the same as mentioned in the legend of Fig. 6A. Representative images from three biological replicates are shown.

**Fig. S6** Rescue of oxidative stress phenotype by ROS scavengers in *C. trachomatis* L2. (A) IFA of wild-type *Ctr* L2 incubated or not with scavengers, α-Tocopherol (100 µM) and DMTU (10 mM), and treated or not with 500 µM H_2_O_2_ as mentioned in materials and methods. Representative images from three biological replicates are shown. (B) IFU analysis of *Ctr* L2 grown under the same conditions as mentioned in the legend of Fig. 7G. IFUs were calculated as percentage of untreated samples. \*\*\**p*< 0.0001, statistical analysis was performed using ordinary one-way ANOVA. H_2_O_2_ treated sample and samples incubated with scavengers only were compared to the untreated control. Samples treated with H_2_O_2_ and incubated with scavengers were compared with sample treated with H_2_O_2_. Data represent three biological replicates. Only significant differences are noted.

**Fig. S7** Effects of *ahpC* knockdown/overexpression on chlamydial developmental cycle progression. RT-qPCR analysis of late cycle genes (*hctA*, *hctB*, *omcB*, *tsp*, and *glgA*) in (A) complementation, and (B) *ahpC* overexpression strain. Experimental conditions were the same as mentioned in the legend of Fig. 7A, and 2A, respectively. Quantified cDNA was normalized to gDNA, and values were plotted on a log scale. \*\*\**p*< 0.0001, \*\**p*< 0.001, \**p*< 0.01 vs uninduced sample by using two-way ANOVA. Data represent three biological replicates.

**Fig. S8** Effect of penicillin treatment during *ahpC* knockdown. (A) IFA was performed to assess inclusion size and morphology of *ahpC* KD-spec following induction (1 nM aTc) and penicillin treatment (1 U/mL) at 10 hpi. At 24 hpi, cells were fixed with methanol and stained using primary antibodies to major outer membrane protein (MOMP), Cpf1 (dCas12), and DAPI. All images were acquired on Zeiss Axio Imager Z.2 with Apotome2 at 100x magnification. Bars, 2 µm. Representative images of three biological replicates are shown. Transcriptional analysis of (B) *ahpC*, (C) *hctA*, and (D) *hctB* in *ahpC* KD-spec using RT-qPCR using the same conditions as in section (A). RNA samples were harvested at 16 and 24 hpi. Quantified cDNA was normalized to 16SrRNA, and values were plotted on a log scale. \*\*\**p*< 0.0001, \*\**p*< 0.001, \**p*< 0.01 vs uninduced sample by using two-way ANOVA. Data represent three biological replicates.

